# Integration of transcription regulation and functional genomic data reveals lncRNA SNHG6’s role in hematopoietic differentiation and leukemia

**DOI:** 10.1101/2023.11.22.568210

**Authors:** Joshua M. Hazan, Raziel Amador, Tamar Lahav, Yehuda G. Assaraf, Roderic Guigó, Assaf C. Bester

## Abstract

**Background:** Long non-coding RNAs (lncRNAs) are pivotal players in cellular processes, and their unique cell-type specific expression patterns make them attractive biomarkers and therapeutic targets. Yet, the functional roles of most lncRNAs remain enigmatic. To address the need to identify new druggable lncRNAs, we developed a comprehensive approach integrating transcription factor binding data with other genetic features to generate a machine learning model, which we have called INFLAMeR (Identifying Novel Functional LncRNAs with Advanced Machine Learning Resources).

**Methods:** INFLAMeR was trained on high-throughput CRISPR interference (CRISPRi) screens across seven cell lines, and the algorithm was based on 71 genetic features. To validate the predictions, we selected candidate lncRNAs in the K562 leukemia cell line and determined the effect of their knockdown on cell proliferation and chemotherapy drug resistance. We further performed transcriptomic analysis for candidate genes. Based on these findings, we assessed the lncRNA Small Nucleolar RNA Host Gene 6 (*SNHG6*) for its role in myeloid differentiation by incubation with Phorbol 12-myristate 13-acetate (PMA) to induce megakaryocyte differentiation, or with hemin to induce erythrocyte differentiation.

**Results:** The INFLAMeR model successfully reconstituted CRISPRi screening data and predicted functional lncRNAs that were previously overlooked. Intensive cell-based and transcriptomic validation of nearly fifty genes in K562 revealed cell type-specific functionality for 85% of the predicted lncRNAs. Our cell-based and transcriptomic analyses predicted a role for SNHG6 in hematopoiesis and leukemia. Consistent with its predicted role in hematopoietic differentiation, *SNHG6* transcription is regulated by hematopoiesis-associated transcription factors. Knockdown of SNHG6 reduced the proliferation of leukemia cells and sensitized them to differentiation. Treatment of K562 leukemic cells with hemin and PMA, respectively, demonstrated that SNHG6 inhibits red blood cell differentiation but strongly promotes megakaryocyte differentiation. Despite *SNHG6* transcripts showing strong cytoplasmic enrichment, *SNHG6* regulates the expression of hematopoietic genes such as *PPBP* (Pro-Platelet Basic Protein) and *PF4* (Platelet Factor 4).

**Conclusions:** Our approach not only improved the identification and characterization of functional lncRNAs through genomic approaches in a cell type-specific manner, but also identified new lncRNAs with a role in hematopoiesis and leukemia. Such approaches cab be used to identify new targets for precision therapy.

## Background

Long non-coding RNAs (lncRNAs) are RNA transcripts that exceed 200 nucleotides in length and have low or no protein-coding potential [1–3]. The transcription of most lncRNAs is regulated through the same mechanism as that of protein-coding genes (PCGs), involving RNA polymerase II and transcription factors (TFs). Furthermore, lncRNAs share several characteristics with PCGs, including a poly-A tail and gene bodies consisting of exons and introns. They comprise one of the largest groups of non-coding elements in the human genome, with an estimated number of annotated lncRNAs ranging from 15,000–150,000 genes [1–4]. Despite this, lncRNAs are one of the most poorly understood groups of non-coding elements in terms of functional characterization [5,6]. Identifying functional lncRNAs remains a challenge due to experimental limitations and an incomplete understanding of their biology.

Experimentally, the functionality of large groups of lncRNAs can be studied through reverse genetic screenings. Library-based screens using various gene perturbation methods have been performed, including antisense oligonucleotides [7], short hairpin RNA with engineered siRNAs [8], and CRISPR-Cas9 [1,9]. While CRISPR has revolutionized reverse genetics by increasing accuracy for the study of PCGs, Cas9-based knockout is not effective for lncRNAs, which are minimally sequence-dependent. Nevertheless, several library screens have been performed using this method, typically by targeting splice sites [10] or with paired gRNAs designed to remove entire exons, introns, promoters, or transcription start sites (TSSs) from the genome [1,11,12]. Alternative CRISPR-based tools have been developed to modulate gene expression without modifying the genome by using a nuclease-dead Cas9 (dCas9) that binds the target sequence without cleaving the DNA. This can be combined with repressive effector protein domains for CRISPR inhibition (CRISPRi) [13] or with transcription activating domains for CRISPR activation (CRISPRa) [14,15]. Additionally, other CRISPR-based approaches are under development, such as stable CRISPR epigenetic silencing (CRISPR-off) and Cas13, which targets RNA [16]. However, screening approaches are limited to specific cell lines and tend to suffer from low sensitivity and accuracy [1].

Computational methods have emerged as a promising approach for predicting the functionality of lncRNAs. These methods include investigating evolutionary conservation [17] and sequence similarities [18], as well as considering expression, differential expression, or co-expression in specific biological contexts [19]. In recent years, multi-omics data integration has been used to complement these approaches and improve their accuracy [20].

Recent advances in machine learning (ML) techniques show great potential for adaptation to address this complex problem. Supervised ML is used for classifying elements based on highly complex data, high dimensionality, or high volumes of data. Indeed, by integrating multiple sources of information, ML models have already been used to successfully predict complex regulatory elements such as enhancers [21], as well as to identify gene regulatory networks and the functional roles of non-coding RNAs in gene regulation. While ML has successfully distinguished lncRNAs from mRNAs [22] and classified tumors based on lncRNA expression [23], predicting the functionality and sub-classification of lncRNAs remains a challenging task. Identifying meaningful features and reliable training sets is essential for classifying functional lncRNAs in specific biological contexts.

To bridge this gap, we developed an ML model known as INFLAMeR (Identifying Novel Functional LncRNAs with Advanced Machine Learning Resources). The model was based on 143 genetic features that focus on cell type-specific regulatory mechanisms and was trained on previous CRISPRi screens [24–26]. INFLAMeR successfully replicated the CRISPRi screening results and identified additional lncRNAs with high prediction scores. Through cell-based and transcriptomics analyses, we tested the true positive and true negative accuracy of INFLAMeR’s predictions for more than forty lncRNAs *in vitro*. Experimental validation based on the INFLAMeR scores demonstrated high predictive accuracy. Of all the lncRNAs with high INFLAMeR scores, 85% displayed significant phenotypic impacts upon KD, whereas lncRNAs with low INFLAMeR scores did not show any effects in the same assays. We further characterized the lncRNA small nucleolar host gene 6 (*SNHG6*) and uncovered its role in hematopoietic differentiation. Overall, our results indicate that INFLAMeR enhances the prediction capacity of functional lncRNAs to a greater extent than large-scale screening assays and can identify lncRNAs for characterization in a cell-type-specific manner.

## Methods

### Developing the ML algorithm

#### ENCODE TF ChIP-seq

We used ENCODE TF ChIP-seq data to determine TF peak height within lncRNA promoters across five cell lines; HEK293T, HeLa, MCF7, K562, and H1-hESC; using 124 TFs.

We downloaded the bigBed narrowPeak files with optimal irreproducible discovery rate thresholded peaks in *hg19* assembly coordinates. We applied a window of [-300; +100] bp surrounding the TSS to obtain lncRNA promoters, as described previously [27]. Next, using *BEDTools* intersect version: 2.27, TFs bigBed, and lncRNA promoters bed files, the TF peak height was obtained. A 10% intersection cutoff between TF ChIP-seq and lncRNA promoter was used.

#### Model training

Stratified 10-fold cross-validation with three different randomizations in each repetition was adopted to train all supervised models, using the *RepeatedStratifiedKFold* class from *scikit-learn* version 0.24.1, with 90% and 10% for training and test, respectively (Supp. Fig. S1g).

#### XGBoost

The *dmlc XGBoost* library (https://xgboost.readthedocs.io/en/latest/index.html) version 1.3.3 was used for implementing the XGBoost model [28]. XGBoost is a type of gradient boosting decision tree method.

As our dataset was unbalanced, the ratio of the minority positive class (hits) versus the majority negative class (not hits) was 1:55; we adopted the XGBoost *scale position weight* parameter to train a cost-sensitive XGBoost classifier for imbalanced data. The default *scale position weight*: 54.81 [sum(majority negative class)/sum(minority positive class)], 100, and 1000 values; in addition to other XGBoost hyper-parameters, were used for grid search. XGBoost was tuned to search for an optimal sensitivity and specificity solution. To tune the hyper-parameters, we adopted the *GridSearchCV scikit-learn* class to improve the performance of the model, using a NVIDIA GPU GeForce RTX-2060 (drivers version 465.31, and CUDA version 11.3). The hyper-parameters tuned for XGBoost and their final values are displayed below (Supp. Fig. S1f):

- Scale position weight: 100
- Learning rate: 0.05
- Max depth: 5
- Regularization lambda: 5.0
- Gamma: 1.0

#### Metrics

To evaluate model performance, we determined the sensitivity (recall), specificity, precision, F1 score, AUROC, AUPRC, Brier score, and Brier skill score, using the test set and the stratified 10-fold cross-validation process described above.

#### Recursive feature elimination (RFE)

Recursive feature elimination (RFE)—removing the least importance features based on SHAP values with stratified cross-validation—was implemented using the *ShapRFECV* class from the *probatus* python module (https://ing-bank.github.io/probatus/index.html), version 1.8.4. The step was one feature per iteration, with sensitivity and specificity as scoring metrics.

#### Model explainability

TreeExplainer from the SHapley Additive eXPlanations (SHAP) framework version 0.39.0 was used to explain the output of our XGBoost model. Global and local explanations were obtained based on 10% of the data not used to train our algorithm. The SHAP framework is based on Shapley values, which is a cooperative game theory concept [29].

#### Computational settings

All the computational analyses for this section were carried out using Linux-based distributions and Python 3.8.5, with computational resources provided by the Centre for Genomic Regulation (CRG), Spain.

The raw features data used to train the ML can be found in Supp. Table S7.

### Cell culture

The following cell lines were used in our experiments: K562 chronic myeloid leukemia cells and the HEK293T human embryonic kidney cells. K562 cells were maintained in RPMI-1640 growth medium containing L-glutamine and 25 mM HEPES pH 7.4 (Gibco, Waltham, MA, USA), supplemented with 10% FCS (Gibco) and 1% penicillin-streptomycin solution (Sartorius, Goettingen, Germany) and passaged every 2–4 days. HEK293T cells were maintained in Dulbecco’s Modified Eagle’s Medium (DMEM; Rhenium, Modiin, Israel) supplemented with 10% FCS, 1% penicillin-streptomycin solution, and 1% L-glutamine (Sartorius) and passaged every 2–4 days.

### Stable transduction of plasmids into K562 cells

For KD by CRISPRi, K562 cells were stably transduced with the pHR-SFFV-dCas9-BFP-KRAB cassette (Addgene plasmid #46911; kindly provided by Stanley Qi & Jonathan Weissman) [30]. Transduction was performed via lentiviral delivery using PolyJet In Vitro Transfection Reagent (SignaGen, Frederick, MD, USA) following the manufacturer’s protocol. Briefly, HEK293T cells were seeded 18–24 h before transfection to obtain a cell confluency of ∼90% at the time of transfection. The cells were transfected with the plasmid of interest, as well as the VSV-G (Addgene plasmid #8454) and psPAX2 (Addgene plasmid #12260) lentiviral packaging plasmids. Lentivirus was harvested 48–72 h after transfection. Lentiviral transduction was performed by incubating K562 cells with DMEM containing the harvested lentivirus for 24 h; next, the media was replaced with fresh RPMI-1640 growth medium, and cells were allowed to recover for 48– 72 h. Finally, cells stably expressing BFP were selected by fluorescence-activated cell sorting (FACS) using the BD FACSAria-IIIu (BD Biosciences, Franklin Lakes, NJ, USA) to obtain the K562-CRISPRi stable cell line.

The gRNAs used for KD were obtained from a previous lncRNA-specific gRNA library [24]. The gRNAs were cloned into either pCRISPRia-v2 (kindly provided by Jonathan Weissman; Addgene plasmid #84832) [31] or pSB700 expressing blasticidin resistance (pSB700-Blast; kindly provided by George Church; Addgene plasmid #64046) [32]. To confirm successful incorporation of the insert, *E. coli* DH5α colonies transformed with the cloned plasmids by heat-shock were subjected to colony PCR using three primers: two primers surrounding the insert site and a third primer overlapping the sequence that is removed during cloning. The PCR products were visualized after 2% agarose gel electrophoresis at 160 V for 30 min. The backbone plasmid with no insert showed two bands of 800 bp and 500 bp for pCRISPRia-v2, or 600 bp and 150 bp for pSB700; plasmids with successful incorporation of the insert showed the larger band only. The primers used to confirm successful ligation can be found in Supp. Table S8.

The gRNAs were stably transduced into K562-CRISPRi cells via lentiviral transduction as described above. For each lncRNA, two gRNAs targeting the same gene were simultaneously transduced. The first gRNA was cloned into pCRISPRia-v2 (expressing tagBFP and puromycin resistance), and the second gRNA was cloned into pSB700-Blast. After incubating the cells with growth medium containing the harvested lentivirus for 24 h, the medium was replaced with fresh RPMI-1640 growth medium supplemented with 1 μg/ml puromycin and 10 μg/ml blasticidin every 48–72 h for 7–10 days. Additionally, cells were transduced with plasmids containing non-targeting gRNAs and subjected to antibiotic selection or selected by FACS. The sequences of the gRNAs used can be found in Supp. Table S9.

### RNA isolation and RT-qPCR

Following stable cell transduction with the indicated gRNAs, RNA extraction was performed using TRI reagent according to standard protocols. RNAs were reverse transcribed to cDNAs using the Quantabio qScript cDNA synthesis kit (QIAGEN, Beverley, MA, USA). The cDNAs were then used for qPCR using qPCRBIO SyGreen Blue Mix Lo-ROX (PCR Biosystems, London, UK), and expression was measured relative to that of cells transduced with a non-targeting sgRNA. Changes in gene expression were calculated using the standard 2^-ΔΔCt^ method and normalized to GAPDH and PGK1 [33]. The primers used for qPCR are listed in Supp. Table S8.

### Two-color competitive cell growth assay

To determine the effect of KD on cell growth and proliferation, each sample was subjected to a two-color CCG assay [24,34] as follows. K562-CRISPRi cells transduced with a non-targeting sgRNA (expressing GFP) were co-seeded at 2×10^5^ cells/ml in the same well as cells transduced with sgRNAs targeting the indicated gene (expressing BFP) at 2×10^5^ cells/ml in 2 ml RPMI-1640 growth medium (day 0). The percentage of GFP- and BFP-expressing cells was determined every 2–4 days by flow cytometry for 14 days. Relative cell growth and proliferation was calculated as the proportion of BFP-expressing cells normalized to that at day 0.

### Drug sensitivity assay

Daunorubicin hydrochloride (DNR) was purchased from Merck (Darmstadt, Germany) and dissolved in dimethyl sulfoxide (DMSO). To determine the optimal concentration of DNR for the drug sensitivity assay, K562 cells with a non-targeting sgRNA were seeded at 2×10^4^ cells/ml in 100 µl RPMI-1640 growth medium supplemented with 1 nM–30 µM DNR for 72 h with n = 2 biological replicates for each concentration [35]. The fraction of surviving cells was determined by flow cytometry and normalized to that of cells incubated in DNR-free growth medium for the same period. A concentration of 1 µM DNR was used for the drug sensitivity assay.

Each sample was seeded at 2×10^4^ cells/ml in 100 µl RPMI-1640 growth medium supplemented with 1 µM DNR for 72 h with n = 3 biological replicates. The proportion of surviving cells was measured by flow cytometry and normalized to that of cells from the same sample incubated without DNR for the same period.

### RNA Sequencing

RNA sequencing was performed for K562 cells with sgRNAs targeting the following lncRNAs: non-targeting sgRNA 10010, non-targeting sgRNA 10057, *AC005307.3*, *AP006222.2*, *CHD1-DT*, *LINC00221*, *MIR4435-1HG*, *RP11-109M17.2*, *RP11-307E17.8*, *RP11-706O15.3*, and *SNHG6*. Total RNA was extracted from n = 3–4 biological replicates per sample using the PureLink RNA Mini Kit (Invitrogen, Waltham, MA, USA), and sample quality was evaluated using a TapeStation. RNA sequencing was performed using CEL-Seq2 [36], a multiplexed RNA-Seq approach that can be used for pooled bulk RNA Sequencing. Raw reads were processed with the Galaxy web platform, using the public server at usegalaxy.org [37]. Briefly, the raw files were converted to FASTQ format using FASTQ Groomer [38], alignment was performed using HISAT2 [39], and counts were generated using featureCounts [40].

Differential gene expression analysis was performed using the NOISeq package in R [41,42]. Since the samples were sequenced in two batches, a strong batch effect was observed between the two runs (Supp. Fig. S9a); additionally, we observed a large difference between the nine KD samples and the two non-targeting control samples (Supp. Fig. S9a). Batch effect correction was performed using NOISeq (Supp. Fig. S9b). After correcting for batch effect, we still observed a high proportion of overlapping differentially expressed genes (DEGs) for each hit versus the controls, indicating that many identified DEGs were likely due to batch effect; therefore, we discarded the control samples and instead analyzed each hit compared to the other eight samples to obtain a more accurate indication of the DEGs in each sample (Fig. 3b). Strongly DEGs were defined as genes with |Log2 Fold Change| > 0.7 and probability > 0.75. Gene Ontology (GO) analysis was performed for the *SNHG6* KD sample based on DEGs using the Enrichr web platform for GO Biological Process [43–45]. To determine the enrichment of TFs associated with the DEGs, the same genes were analyzed using the ChEA3 web platform [46]. To identify the TFs that bind the *SNHG6* promoter in K562 cells, we downloaded the TF binding data for the [-300; +100] bp region surrounding the TSS from the ENCODE 3 Transcription Factor ChIP-seq Peaks track in the UCSC genome browser for the hg19 reference genome [47–52].

### Megakaryocyte differentiation assay

Phorbol 12-myristate 13-acetate (PMA; Merck, Darmstadt, Germany) was dissolved in DMSO. Cells were seeded at 3×10^5^ cells/ml in 1 ml RPMI-1640 growth medium supplemented with 0.2 nM PMA for 72 h, with fresh growth medium replaced daily. Megakaryocyte differentiation was determined based on CD41/CD61 membrane protein levels [53]. The fraction of CD41/CD61-positive cells was determined after incubation for 24, 48, and 72 h by immunofluorescent staining with a PE-conjugated anti-human CD41/CD61 primary antibody (BioLegend, San Diego, CA, USA) using flow cytometric analysis.

### Erythrocyte differentiation assay

Hemin (Merck, Darmstadt, Germany) was dissolved in DMSO. Cells were seeded at 3×10^5^ cells/ml in 1 ml RPMI-1640 growth medium supplemented with 30 µM hemin for 72 h, with fresh growth medium replaced daily. Erythrocyte differentiation was determined based on glycophorin A (GPA) membrane protein levels [54]. The fraction of GPA-positive cells was determined after incubation for 24, 48, and 72 h by immunofluorescent staining with a PE-conjugated anti-human GPA primary antibody (BioLegend, San Diego, CA, USA) using flow cytometric analysis.

### Statistical analysis

Statistical analyses were performed using the rstatix package in R. Values are given as the mean ± SD. Significance was evaluated using a two-tailed Student’s *t*-test with Benjamini-Hochberg correction unless otherwise stated. A value of *p* < 0.05 was considered statistically significant.

## Results

### An XGBoost classifier to uncover the function of lncRNAs in cell growth

The human genome encodes more than 15,000 lncRNAs, which are a highly diverse group of genes. While some lncRNAs encode their function in their sequence, others act as DNA regulatory elements, such as enhancers [55]. Furthermore, it remains unclear how many lncRNAs are functional compared to those that are expressed as transcriptional noise. To better understand the biology of lncRNAs, it is necessary to classify them according to their function.

Over the last several years, ML has been exploited to identify complex relationships between biological properties and predict functionality in various contexts [56]. Herein we developed a binary classification model to identify new functional lncRNAs based upon genetic features, which were trained using known functional lncRNAs (Fig. 1a, upper panel).

**Figure 1.**
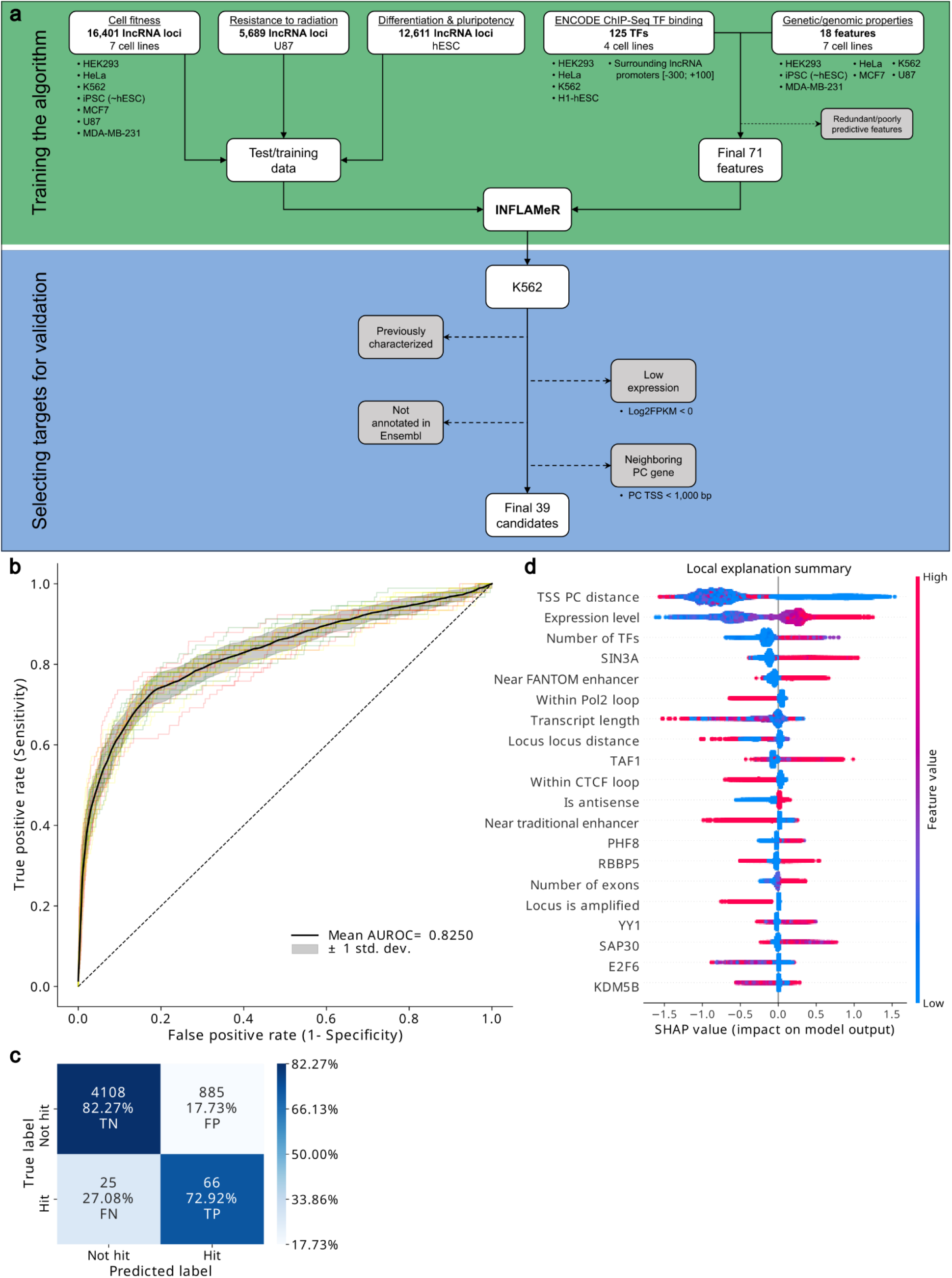
Building a machine learning algorithm for the accurate prediction of novel functional lncRNAs. (a) Workflow for designing the INFLAMeR (Identifying Novel Functional LncRNAs with Advanced Machine learning Resources) algorithm and selecting targets. Upper: The algorithm was trained on previous high-throughput pooled CRISPR interference (CRISPRi) screening data from three previous high-throughput CRISPRi screens [24–26]; the algorithm was computed based on a total of 71 features comprising ENCODE ChIP-Seq transcription factor (TF) binding data for the regions surrounding lncRNA promoters [50–52] and genomic features. Lower: targets predicted to be functional by INFLAMeR in the K562 cell line (INFLAMeR score > 0.5) were selected for validation after excluding lncRNAs that were functionally characterized in previous studies, those with low expression, those not annotated in the Ensembl database, and those neighboring a protein-coding (PC) gene; thirty-nine lncRNAs were selected for validation. (b) INFLAMeR was built using an XGBoost classifier and its performance was calculated using a receiver operating characteristic curve. Black ROC curve shows the mean classifier performance on the test set using three randomized seeds (red, green, and yellow curves). (c) Confusion matrix based on the test set. Percentages in the confusion matrix are row-normalized. (d) Local explanation summary of the impact of the top twenty features on the INFLAMeR score. Each dot represents one lncRNA. Red indicates a higher feature value (e.g., larger transcription start site (TSS) protein-coding (PC) distance or higher expression), and blue indicates a lower feature value (e.g., smaller TSS PC distance or lower expression). Higher (red) feature values with a positive SHAP value indicate a positive correlation, and lower (blue) feature values with a positive SHAP value indicate a negative correlation.

To create the ML algorithm, we gathered 143 genetic variables representing different features of lncRNAs (Supp. Table S1). These features included eighteen categorical variables related to lncRNAs and their surrounding genetic regions, such as the transcript length, number of exons, proximity to enhancers, and proximity to PCGs. While these features are important, most do not take into account the cell type-specific functions of lncRNAs. Since most lncRNAs exhibit cell type-specific functionality [1,5], we also included features pertaining to regulation and expression, namely TF binding data from the ENCODE project for lncRNA promoters in four different cell lines for more than 100 TFs [50–52]. We then trained the ML algorithm using a set of functionally validated lncRNAs. Although many human lncRNAs have been demonstrated to be functional in various contexts, such as cell proliferation [24,57], differentiation [26,58], and disease [20,59,60], these lncRNAs have often been studied in different cell types and under varying experimental conditions, resulting in scattered data.

In recent years, high throughput reverse genetic screens have been increasingly utilized to explore the functions of numerous lncRNAs [61]. These screens, which involve pooled perturbations, offer a significant advantage as they allow thousands of genes to be tested in parallel under identical experimental conditions. The resulting scores generated by these screens are statistically based and enable the classification of genes as either hits (functional) or non-hits. While various perturbation methods have been employed to identify functional lncRNAs, CRISPRi has emerged as the most applied technique across multiple cell types and biological contexts [1].

To train our algorithm, we incorporated data from three genetic screens [24–26] that targeted 16,401 lncRNAs across seven cell lines. However, these data are imbalanced since only 9% (n = 1,451) of the lncRNAs were identified as functional. Moreover, perturbation screening of lncRNAs is subject to high false-negative rates due to the inconsistent efficiency of CRISPRi [62] and poor annotation of lncRNA TSSs. This dataset imbalance may introduce a bias into the training algorithm, resulting in diminished predictive value.

To develop the ML model, we experimented with three different cost-sensitive classifiers: logistic regression, balanced random forest, and extreme gradient boosting (XGBoost). We assessed the performance of the model using a range of metrics, including area under the receiver operating characteristic curve (AUROC), sensitivity, specificity, F1-score, precision, and Brier score (Supp. Table S2). Our results show that the mean AUROC for logistic regression was 0.778, which was lower than that for both XGBoost and balanced random forest (mean AUROC values: 0.8236 and 0.8335, respectively; Supp. Fig. S1a).

To determine which ML algorithm including XGBoost or balanced random forest, would be better suited for our desired balance of sensitivity and specificity, we determined the percentages of true positive and true negative results for each method (Supp. Fig. S1b). XGBoost had a somewhat higher specificity percentage than balanced random forest (82.24% vs. 80.84%). On average, both XGBoost and balanced random forest produced a similar number of true positive cases (66 and 69 cases, respectively). Additionally, XGBoost possessed two advantages over balanced random forest. Firstly, in terms of metrics, XGBoost had higher F1-score and precision outcomes (0.1264 and 0.0693, respectively) than balanced random forest (0.1240 and 0.0675). Secondly, in terms of training time, XGBoost had a shorter training time than balanced random forest, which enabled us to experiment and explore more training settings. Therefore, we selected XGBoost for our ML model.

To address the class imbalance problem in our dataset, we applied random under-sampling of non-hits with and without replacement methodologies as preprocessing before training the XGBoost model. We experimented with different sampling strategies to find the optimal ratio of non-hits to hits. The sampling strategies were as follows: 3%, 4%, 5%, 10%, 20%, 30%, 40%, and 50% without and with replacement (Supp. Fig. S1c and d, respectively). We evaluated the performance of each strategy using various metrics such as accuracy, precision, recall, F1-score, and AUROC. We obtained a superior performance for the 50% sampling strategy (1,822 non-hits and 911 hits), both without and with replacement (Supp. Table S3 and S4, respectively). However, we also explored a cost-sensitive approach that assigns different weights to the classes based on their proportion within the dataset. We compared the cost-sensitive XGBoost model with the under-sampled XGBoost models and found that it exhibited superior performance compared to under-sampling with 50% in terms of the AUROC and sensitivity values. Thus, cost-sensitive XGBoost was selected as our ML model.

We used Shapley (SHAP) values [29] to assess the importance of each feature in our XGBoost model for predicting lncRNA function. Out of the 143 features, 30 had no predictive value (SHAP ≈ 0) and were hence excluded from further analysis. Although combining multiple features can enhance the predictive power of the model, it can also introduce high-dimensional redundancy and increase the training time and performance bias [63,64]. Therefore, we applied a recursive feature elimination (RFE) method based on SHAP values to select the optimal subset of features that balanced sensitivity and specificity on the test set. Based on RFE, the best performance was obtained with 71 features (Supp. Fig. S1e).

We then evaluated our cost-sensitive XGBoost model with RFE using a triple-repeated 10-fold cross-validation with stratified sampling. The hyper-parameters of our gradient-boosted tree classifier were as follows: initial guess of 0.5, gamma of 1.0, gain as importance, learning rate of 0.05, residual-trees with 5 depth levels and 28 leaves (Supp. Fig. S1f), 100 residual-trees, random seed of 0, regularization of lambda of 5.0, and a cost ratio of 100 for misclassifying a hit versus a non-hit.

Using 71 features, the cost-sensitive XGBoost method showed better performance than previous methods. In fact, all metrics were higher for the 71-feature model compared to the 143-feature model; sensitivity and AUROC were increased by 0.05 and 0.002, respectively (Supp. Table S5). We also compared the final XGBoost model to our previous approach using balanced random forest. The XGBoost model with RFE displayed higher mean values of specificity, F1-score, and precision (0.8227 vs. 0.8084, 0.1275 vs. 0.1240, 0.0698 vs. 0.0675, respectively) than the balanced random forest model (Supp. Table S5).

To estimate the probability of hits among all lncRNAs, we applied our trained model to classify the 50,847 transcripts in our dataset. Transcripts with a predicted probability greater than 0.5 were defined as predicted hits. To evaluate the performance of our model, we used a receiver operating characteristic (ROC) curve (Fig. 1b) and a confusion matrix (Fig. 1c). Our model achieved a mean AUROC of 0.8250 ± 0.01 (with minimum and maximum values of 0.78 and 0.87, respectively; see Supp. Table S6 for all AUROC values), with a true positive rate of 72.92% and a true negative rate of 82.27%. Notably, the predicted hit and non-hit values were well balanced. The average F1-score, precision, and Brier scores were 0.1275, 0.0698, and 0.1634, respectively.

Statistical analysis of the impact of individual features on the score confirmed the importance of lncRNA-PCG distance (Fig. 1d). Interestingly, the impact of the distance on the prediction score was not linear; rather, we observed a decrease in the score in a stepwise manner, with the SHAP values sharply decreasing around 1,000 and 2,000 bases from the nearest PCG. However, the distance between promoters did not further affect the score beyond 2,000 bases (Supp. Fig. S1h). Further important features were those affecting transcription, such as transcription level (defined as log FPKM), the number of TFs binding to the promoter, and the binding of specific TFs including SIN3A (Fig. 1d). For example, the score of the lncRNA *SNHG6* was decreased, owing to its distance from a PCG and proximity to a traditional enhancer; however, it was increased based on its high expression and SIN3A binding at its promoter (Supp. Fig. S1i).

Interestingly, although transcription was tightly correlated with lncRNA functionality (Wilcoxon test, *p* = 4.20×10^-222^), the presence of general TFs such as TAF1 and TAF7 did not contribute to the prediction model. These findings suggest that functional lncRNAs may be regulated by a subset of specialized factors.

Our findings indicate that lncRNAs have distinctive features that are closely linked to their functionality. These attributes can be influenced by various factors, such as their genomic location (for example, their proximity to protein-coding genes) and by cell type-specific mechanisms that are intricately regulated by TFs. By identifying these characteristics, one can gain a deeper understanding of the complex roles that lncRNAs may play in regulating genes and cellular processes.

### Validation of INFLAMeR’s predictions and identification of functional lncRNAs

To assess the accuracy of INFLAMeR, we selected a subset of lncRNAs for CRISPRi-mediated knockdown (KD) in K562 cells based on their INFLAMeR score and according to several cutoff parameters (Fig. 1a, lower panel). We excluded the following lncRNAs for validation: those identified in previous screens to be functional in K562 cells; those with low expression (Log2 FPKM < 0) in K562 cells; those which are not annotated in the Ensembl database; and those located within 1 kb of the nearest PCG TSS, as transcription can be initiated from divergent promoters. We included thirty-nine lncRNAs predicted to be functional in K562 cells, with an INFLAMeR score ≥ 0.5. Furthermore, we randomly selected seven lncRNAs that did not meet the INFLAMeR threshold for functional prediction (INFLAMeR score < 0.4) to validate their non-functionality using the same parameters mentioned above. This was done in order to exclude the possibility that the selected lncRNAs were functional by chance.

Predicting CRISPRi-mediated KD efficiency can be challenging, especially for lncRNAs with multiple splice variants. To enhance KD efficiency, we pooled sgRNAs for each target, with each sample containing two sgRNAs targeting the same lncRNA [65]. We confirmed efficient KD, defined as relative expression ≤ 0.5 by qPCR, for 80% of the targeted lncRNAs (n = 37/46) (Supp. Fig. S2).

To determine whether the validated lncRNAs affected cell proliferation upon KD, we conducted a two-color competitive cell growth (CCG) assay for each sample (Fig. 2a); briefly, cells with KD of each lncRNA (or transduced with a non-targeting sgRNA) expressing blue-fluorescent protein (BFP) were mixed at a 1:1 ratio with cells transduced with a non-targeting sgRNA expressing green-fluorescent protein (GFP), and the relative proportion of BFP-positive cells was measured over 14 days by flow cytometry to determine the impact of KD on cell proliferation. We observed a significant reduction in the proportion of BFP-positive cells (*p* < 0.05) for 74% of the samples predicted to be functional (n = 29/39), while none of the seven lncRNAs predicted to be non-functional had an impact on cell proliferation (Fig. 2b).

**Figure 2.**
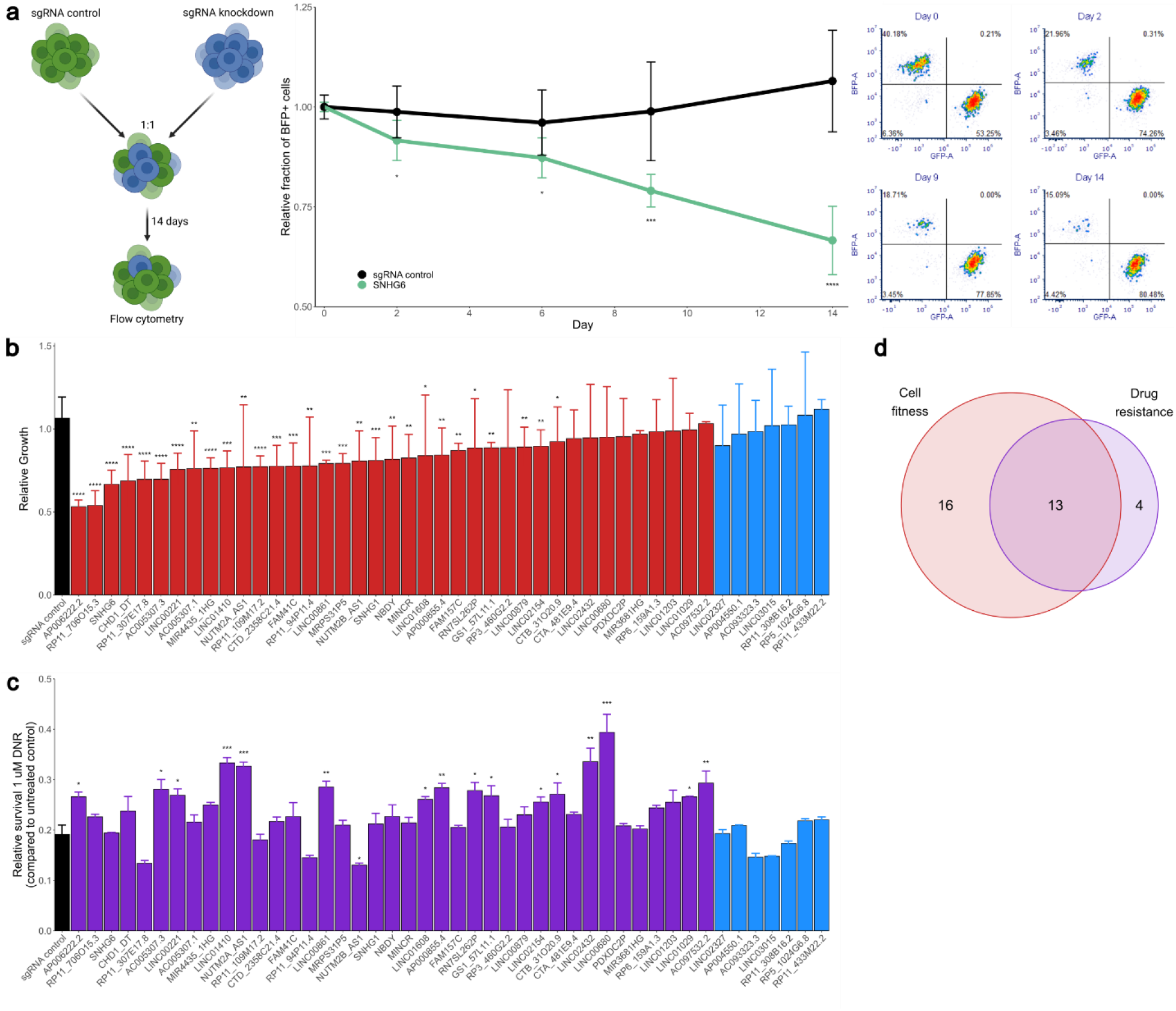
Most lncRNAs predicted to be functional showed a functional effect upon knockdown (KD). (a) Two-color competitive cell growth (CCG) assay. Left: K562 cells stably transduced with sgRNAs targeting a given lncRNA (or non-targeting sgRNA control)—including a blue fluorescent protein (BFP) marker gene—were mixed at a 1:1 ratio with cells transduced with sgRNA control—including a green fluorescent protein (GFP) marker gene—and the fraction of BFP-expressing cells was tracked over 14 days by flow cytometry. The effect of KD on cell proliferation was expressed as the fraction of BFP-expressing cells relative to that at day 0. Middle: a representative example of the change in the relative fraction of BFP-expressing cells throughout the experiment, normalized to day 0. Right: Representative examples of the BFP- and GFP-expressing fractions based on flow cytometry throughout the experiment. (b) The relative growth of BFP-expressing cells after transduction with the sgRNA control (black), thirty-nine lncRNAs predicted to be functional by INFLAMeR (red), and seven lncRNAs predicted to be non-functional (blue) at day 14 of the CCG assay. Error bars represent SD from n = 2 biological replicates. \**p* < 0.05, \*\**p* < 0.01, \*\*\**p* < 0.001, \*\*\*\**p* < 0.0001 *vs.* sgRNA control using a one-tailed *t*-test. (c) Relative cell survival after 72 h incubation with 1 µM daunorubicin (DNR)— relative to that in cells grown in drug-free medium—for sgRNA control (black), thirty-nine lncRNAs predicted to be functional (purple), and seven lncRNAs predicted to be non-functional (blue). Error bars represent SD from n = 3 biological replicates. \**p* < 0.05, \*\**p* < 0.01, \*\*\**p* < 0.001, *vs.* sgRNA control using a two-tailed *t*-test with Bonferroni correction. (d) Of the thirty-nine predicted lncRNAs, many of those with an effect on cell proliferation (red) also affected resistance to DNR (purple) upon KD.

Since lncRNAs often function in *cis*, we investigated whether the validated genes were adjacent to PCGs considered essential based on dependency scores obtained using the DepMap portal, which characterized essential genes based on gene perturbation across more than 1,000 cancer cell lines [66]. We found no correlation between the INFLAMeR score of the thirty-nine predicted lncRNAs and the essentiality of their neighboring PCGs, further supporting the independent function of the lncRNAs (Supp. Fig. S3).

Studying cell proliferation under optimal growth conditions may overlook lncRNAs that function in response to stress cues. Therefore, we incubated each sample with daunorubicin (DNR), an anthracycline chemotherapeutic agent commonly used to treat leukemias [35]; DNR exerts its pharmacologic activity as a DNA topoisomerase II inhibitor. After confirming the optimal concentration of DNR for the growth inhibition assay (Supp. Fig. S4; see methods), each sample was incubated with 1 µM DNR for 72 h to determine whether these lncRNAs affected resistance to DNR. Of the thirty-nine hits, 44% (n = 17/39) exhibited a significant effect on DNR resistance (Fig. 2c), while the seven non-hits did not affect DNR resistance. Of the seventeen lncRNAs affecting DNR resistance, thirteen also showed a significant reduction in cell proliferation from the CCG assay (Fig. 2d).

Thus, 85% (n = 33/39) of the lncRNAs predicted by INFLAMeR to be functional were experimentally validated to have a function in either cell proliferation or anticancer drug resistance. Importantly, most of these lncRNAs have never been functionally characterized until now, and none of them has a known function in K562 cells.

Collectively, these findings demonstrate the utility of INFLAMeR in predicting the functionality of lncRNAs.

### Validated lncRNAs with a strong impact on cell proliferation act in *trans* and affect distinct molecular pathways

To further characterize their mechanism of function, we performed transcriptomic analysis for nine lncRNAs with a strong impact on cell proliferation upon KD. An analysis of the most variable differentially expressed genes (DEGs) for each lncRNA revealed distinct patterns of gene expression for each of the samples (Fig. 3a). This implies that although each lncRNA caused impaired cell proliferation upon KD, this phenotype was presumably achieved via a distinct mechanism for each lncRNA. Similarly, principal component analysis showed that most of the samples were distinctly clustered (Fig. 3b). We also observed that all nine lncRNAs in our analysis affected the expression of genes across the genome, implying that they function via a *trans* regulatory mechanism (Fig. 4a, Supp. Fig. S5). This is in contrast to the known *cis* regulatory function of most characterized lncRNAs. Additionally, for most of the hits, we observed a higher proportion of downregulated DEGs than of upregulated DEGs (Fig. 4b, Supp. Fig. S6). This is consistent with the gene regulatory function of known lncRNAs, many of which are known to act as activators of transcription [2]. For most of the samples, we observed a relatively small number of highly DEGs (between 70 and 300 DEGs—defined as |Log2 Fold Change| > 0.7 and probability > 0.75). Among the nine genes we studied here, *SNHG6* stood out as an interesting candidate.

**Figure 3.**
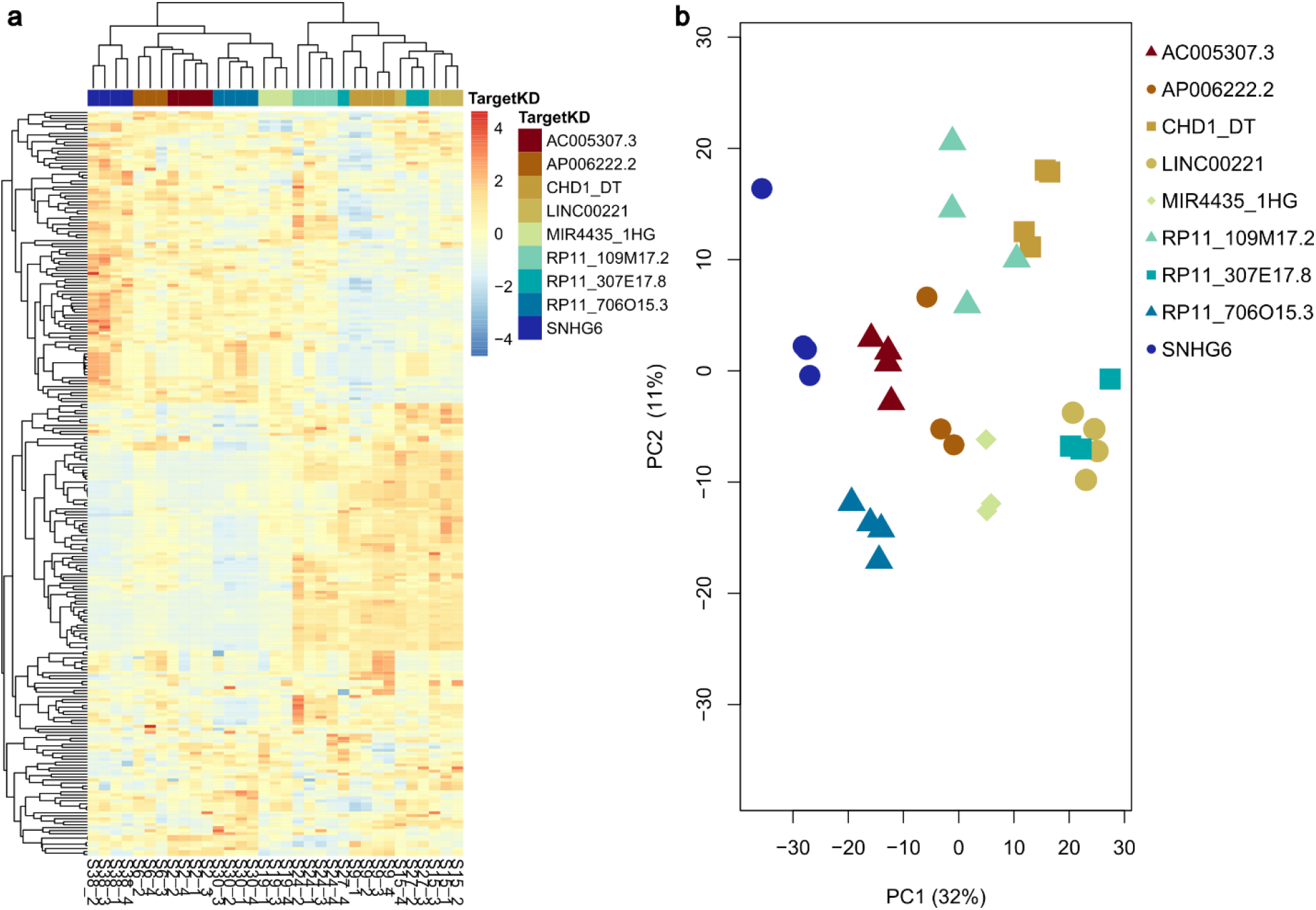
Transcriptome analysis for nine validated lncRNAs. (a) Nine of the validated lncRNAs were subjected to RNA-Seq, followed by differential gene expression analysis. Most of the lncRNAs affected a unique subset of genes upon knockdown (KD). (b) Principal component analysis was performed for the top variable genes in each sample to confirm that each lncRNA affected a distinct group of genes upon KD.

**Figure 4.**
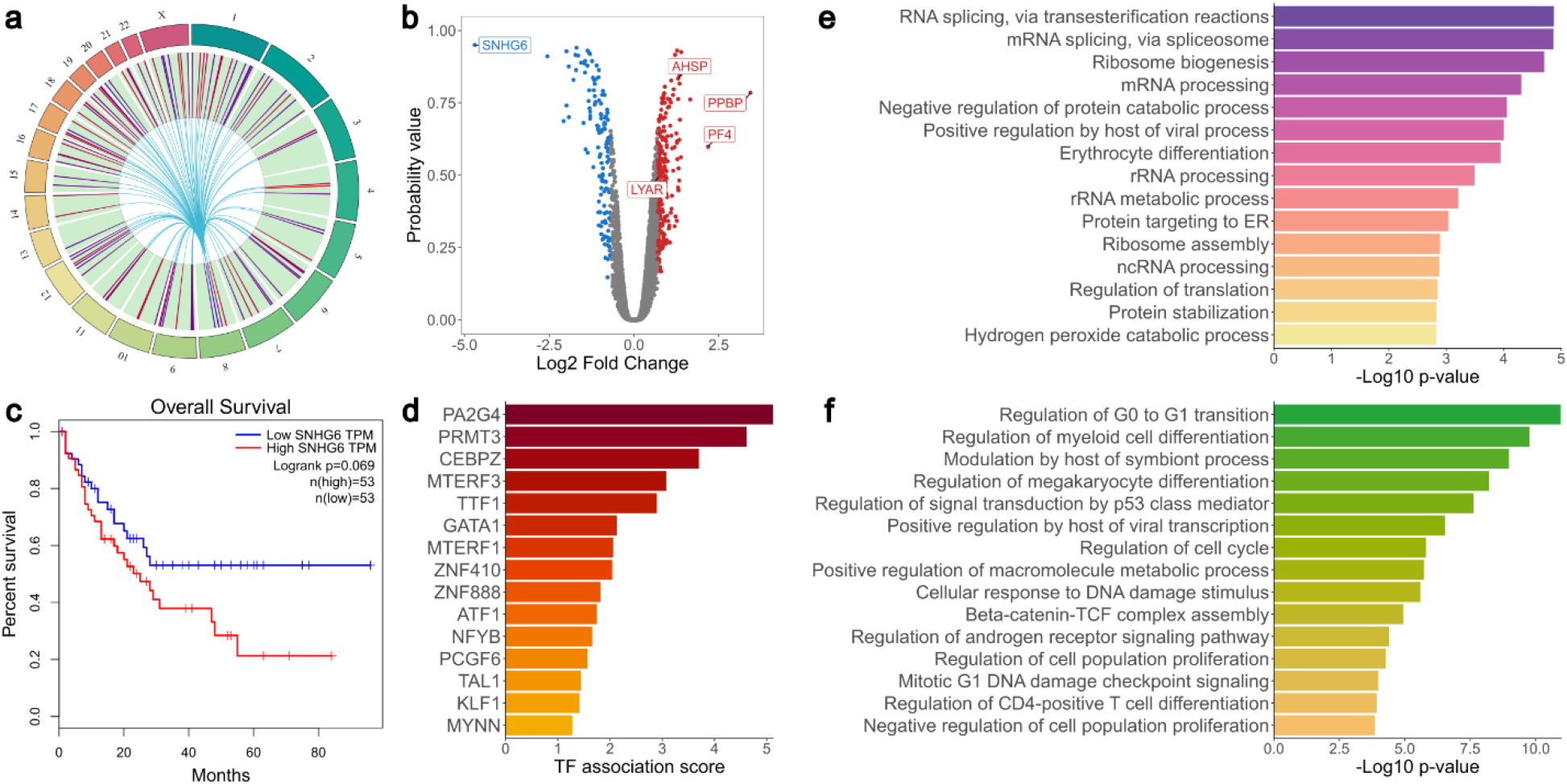
Small nucleolar host gene 6 (*SNHG6*) acts as a regulator of hematopoiesis. (a) *SNHG6* KD affects the expression of genes across the genome, implying that it has a *trans* regulatory impact. Blue represents downregulation, red represents upregulation. (b) *SNHG6* KD led to the upregulation of the platelet-associated genes PPBP and PF4, and the erythrocyte-associated genes AHSP and LYAR. (c) High *SNHG6* expression is weakly associated with reduced survival in LAML. (d) Transcriptions factors (TFs) strongly associated with the differentially expressed genes (DEGs) upon *SNHG6* KD include hematopoiesis-associated TFs such as CEBPZ, GATA1, TAL1, and KLF1. (e) Gene Ontology (GO) analysis of the DEGs indicates a strong correlation with erythrocyte differentiation. (f) GO analysis of the TFs known to bind *SNHG6* in K562 cells shows a strong association with myeloid differentiation-and hematopoiesis-related pathways.

### *SNHG6* regulates hematopoiesis in K562 cells

*SNHG6* is a lncRNA with a known impact in several cancer types [67–73], and is weakly associated with a poor prognosis in acute myeloid leukemia (Fig. 4c).

The KD of *SNHG6* led to the differential expression of 304 genes, including 126 protein-coding genes and 103 lncRNAs. The identified DEGs were widespread across the entire genome (Fig. 4a), suggesting a *trans* regulatory role for *SNHG6*; this was consistent with its strong cytoplasmic localization in K562 cells (Supp. Fig. S7).

Interestingly, *SNHG6* KD led to the upregulation of genes involved in myeloid differentiation to both erythrocyte and megakaryocyte lineages. Following *SNHG6* KD, the most strongly up-regulated genes included the erythrocyte-associated α-hemoglobin stimulating protein (*AHSP*) and Ly1-antibody reactive (*LYAR*); and the megakaryocyte-associated pro-platelet basic protein (*PPBP*) and platelet factor 4 (*PF4*) (Fig. 4b). Using ChEA3, a tool for identifying the TFs most strongly associated with the DEGs [46], we observed a strong correlation with several TFs associated with hematopoietic differentiation, including CEBPZ [74], KLF1, TAL1, and GATA1 [75] (Fig. 4d). Consistent with the erythroleukemia origin of K562 cells, GO analysis using Enrichr [43–45] showed strong enrichment for erythrocyte differentiation (Fig. 4e).

We also performed GO analysis of the TFs targeting the *SNHG6* promoter. Expectedly, the most strongly enriched pathways were associated with transcription (Supp. Fig. S8). Surprisingly, however, there were several highly enriched pathways relating to hematopoietic and myeloid differentiation (Fig. 4f). These findings are in line with recent reports regarding the role of *SNHG6* in hematopoiesis [69,76].

### *SNHG6* regulates hematopoietic differentiation in K562 cells

To corroborate these findings, we assessed the differentiation potential of *SNHG6*-KD cells.

To stimulate erythrocyte differentiation, *SNHG6*-KD and control cells were incubated with hemin, and erythrocyte status was confirmed based on the levels of the surface marker glycophorin A (GPA) after 24, 48, and 72 h (Fig. 5a). Starting at 24 h, we found a significant increase in GPA levels in *SNHG6*-KD cells compared to that in sgRNA control cells (Fig. 5b), indicating an increased affinity for erythrocyte differentiation.

**Figure 5.**
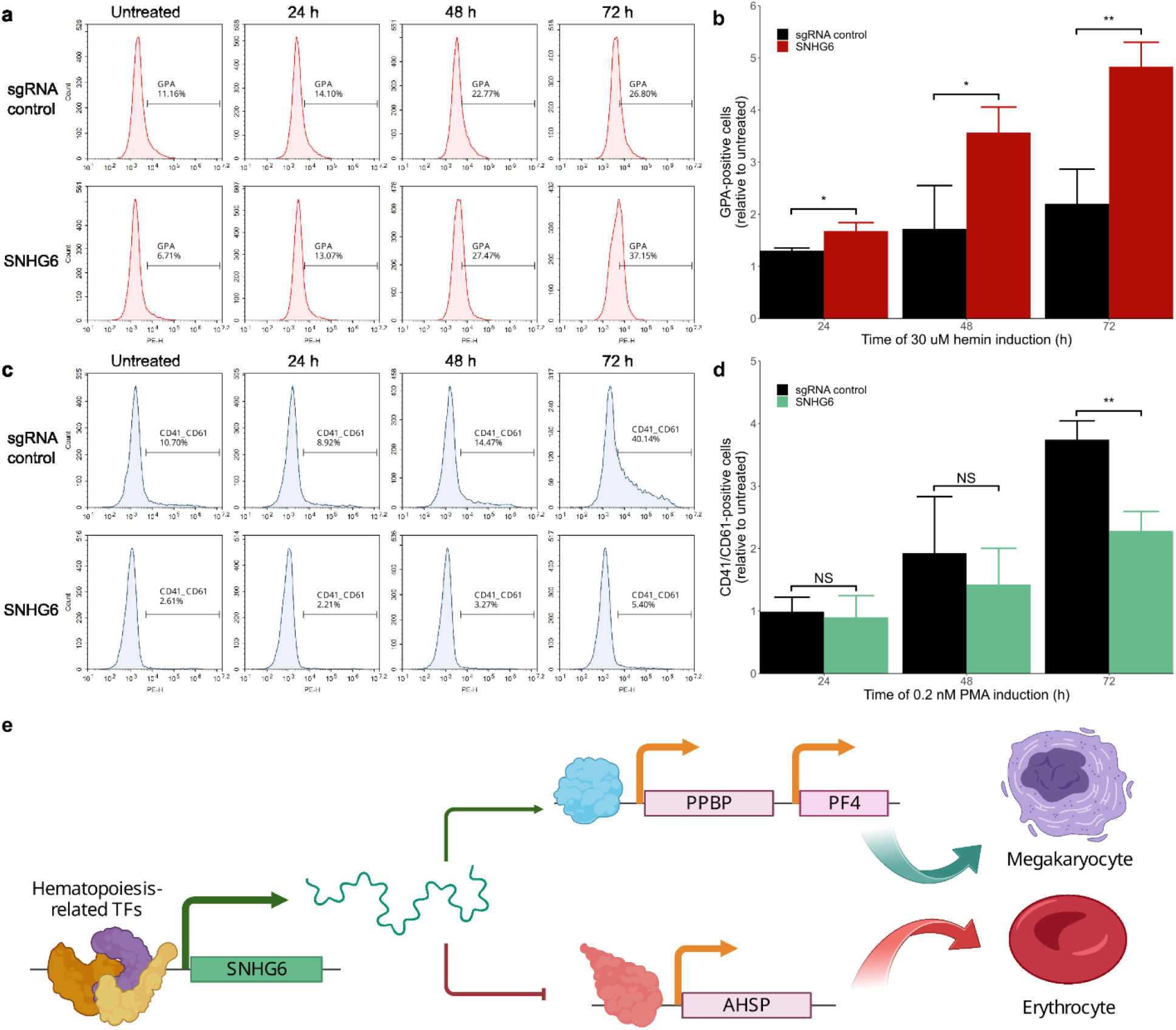
*SNHG6* KD promotes erythrocyte differentiation and prevents megakaryocyte differentiation in K562 cells. (a) Erythrocyte differentiation was assessed based on glycophorin A (GPA) levels using immunostaining with flow cytometry in cells incubated with 30 µM hemin for 72 h. (b) GPA levels were normalized to basal levels in each sample to quantify the degree of erythrocyte differentiation. *SNHG6* KD promoted erythrocyte differentiation. Error bars represent the SD from n = 3 biological replicates. * *p* < 0.05, ** *p* < 0.01 vs. sgRNA control. (c) Megakaryocyte differentiation was measured based on CD41/CD61 levels using immunostaining with flow cytometry in cells incubated with 0.2 nM PMA for 72 h. The basal levels of CD41/CD61 in *SNHG6*-KD cells were substantially lower than in sgRNA control cells. (d) CD41/CD61 levels were normalized to basal levels in each sample to quantify the degree of megakaryocyte differentiation. *SNHG6* KD significantly reduced megakaryocyte differentiation after 72 h. Error bars represent the SD from n = 3 biological replicates. ** *p* < 0.01 vs. sgRNA control. (e) Proposed model for the mechanism of function of *SNHG6*.

To stimulate megakaryocyte differentiation, cells were incubated with phorbol 12-myristate 13-acetate (PMA), and megakaryocyte status was confirmed based on the levels of the CD41/CD61 surface marker after 24, 48, and 72 h. We found that basal CD41/CD61 levels were decreased in *SNHG6*-KD cells compared to those in control cells (Fig. 5c). After 72 h, CD41/CD61 levels were significantly lower in *SNHG6*-KD cells than in sgRNA control cells (Fig. 5d), indicating a decreased stimulation of megakaryocyte differentiation.

Together, our findings show that *SNHG6* KD promotes erythrocyte differentiation and inhibits megakaryocyte differentiation in K562 cells, indicating that *SNHG6* acts as a regulator of hematopoietic differentiation (Fig. 5e).

## Discussion

lncRNAs represent a large and heterogenous group of genes. To enhance our understanding of their biological roles, there is a need to identify subgroups and cluster them accordingly. To this end, the comprehensive characterization of functional lncRNAs is critical. However, this task has proved to be challenging not only due to their diverse functional mechanisms, but also due to their low expression and cell type-specific functions [1,5].

Here, we present INFLAMeR, an ML-based tool for the classification and prediction of functional lncRNAs. INFLAMeR uses both constant and variable genomic features that allow for the prediction in a cell type-specific manner. The variability in lncRNA expression between cell types and cellular conditions indicates that their expression is tightly regulated. Therefore, INFLAMeR was built based on a large set of TFs occupying lncRNA promoters. Indeed, our current analysis of the contribution of different features based on SHAP values indicated that TFs are important contributors for lncRNA classification. Interestingly, functional prediction was correlated with an increased number of TFs; this was the third most strongly contributing feature. This may indicate that the combinatorial effect of multiple factors is critical for their function, rather than the role of a specific TF.

Surprisingly, while the distance between lncRNAs and the closest PCG was the most strongly contributing feature for classification, we did not find a correlation between lncRNAs affecting cell proliferation and the dependency scores of the PCGs neighboring each lncRNA (Supp. Fig. S3). Furthermore, although *cis* regulation is well documented as a mechanism of function for many lncRNAs [77], our transcriptomic analysis following KD of validated lncRNAs did not identify significant changes in the expression of most neighboring PCGs. This suggests that INFLAMeR enriched for *trans* regulating lncRNAs.

Our use of INFLAMeR reveals a large number of false negative results in pooled CRISPRi perturbation screens. We can point out two contributing reasons. Firstly, previous CRISPRi screens transduced only one sgRNA in each cell, which often results in low KD efficiency. Using two sgRNAs targeting the same gene, we achieved a much stronger KD, which allowed us to reveal the function of the targeted lncRNAs. Secondly, in most cases, the overall effect of lncRNA KD on cell proliferation or anticancer drug resistance is relatively mild, as can be expected for regulatory genes. This may result in a low signal-to-noise ratio in a pooled screen. Hence, the improvement of KD efficiency together with post-screening analysis by algorithms such as INFLAMeR may increase the selectivity of perturbation screens and reveal many more functional lncRNAs.

Importantly, intensive validation showed a clear functionality for 85% of lncRNAs displaying a high INFAMeR score, but no functionality for genes with a low score. This confirms the high accuracy of our predictions. Further transcriptomics characterization of the top validated lncRNAs showed that while the number of DEGs was relatively small, each lncRNA affected a different subset of genes. Interestingly, in several cases, lncRNA KD resulted in the coordinated expression of genes in the same locus. However, in the vast majority of cases, the target genes were not neighboring the lncRNA, indicating the *trans*-genomic changes induced by the lncRNAs.

Our analysis identified *SNHG6* as an interesting functional lncRNA. We found that *SNHG6* KD affected cell proliferation, but not response to a *bona fide* cytotoxic agent. These findings are consistent with our transcriptome analysis, which revealed that *SNHG6* regulates hematopoietic differentiation. Both erythrocyte-specific genes such as hemoglobin subunits, as well as platelet-specific genes such as *PPBP* and *PF4*, were differentially expressed following *SNHG6* KD. This KD further caused the erythroleukemic cell line K562 to be susceptible to erythrocyte differentiation by hemin, but resistant to megakaryocyte differentiation. Further studies are warranted to determine whether the observed function of *SNHG6* was caused by the lncRNA itself, or by its contained snoRNA. Additionally, there is recent evidence that *SNHG6* encodes for a small peptide that may be functional [78].

## Conclusion

Overall, we show that INFLAMeR can be trained to readily identify functional lncRNAs in diverse cells and tissue types, as well as under distinct biological cues and contexts. Furthermore, INFLAMeR can be used to augment the sensitivity and specificity of large-scale perturbation screens by constructing more efficient sgRNA libraries.

## Declarations

### Ethics approval and consent to participate

Not applicable.

### Consent for publication

Not applicable.

### Availability of data and materials

The data underlying this article are available in Zenodo, at https://doi.org/10.5281/zenodo.8114662.

## Funding

This work was funded by the Israeli Science Foundation (ISF; grant no. 2228/19) and the Israel Cancer Association (ICA; grant no. 20230069). R.A. was a predoctoral fellow of the CONACYT “Becas al Extranjero” Program of Mexico.

### Author contributions

A.C.B. and J.M.H. initiated the project, designed the experiments, and analyzed the data. J.M.H. performed the experiments and bioinformatics analysis. R.G., R.A., and T.L. designed and implemented the INFLAMeR algorithm. Y.G.A. designed the drug resistance assay. A.C.B., J.M.H., and R.A. wrote the manuscript with input from all the authors. A.C.B. supervised the project.

## Acknowledgements

Not applicable.

## Abbreviations

lncRNA: long non-coding RNA
CRISPRi: CRISPR interference
INFLAMeR: Identifying Novel Functional LncRNAs with Advanced Machine Learning Resources
ML: machine learning
KD: knockdown
SNHG6: Small Nucleolar RNA Host Gene 6
PPBP: Pro-Platelet Basic Protein
PF4: Platelet Factor 4
AHSP: alpha hemoglobin stimulating protein
LYAR: Ly1-antibody reactive
TF: transcription factor
PCG: protein-coding gene
TSS: transcription start site
ROC: receiver operating characteristic
AUROC: area under the receiver operating characteristic curve
RFE: recursive feature elimination
CCG: competitive cell growth
BFP: blue fluorescent protein
GFP: green fluorescent protein
DNR: daunorubicin
DEG: differentially expressed gene
GO: gene ontology
GPA: glycophorin A
PMA: phorbol 12-myristate 13-acetate
DMSO: dimethyl sulfoxide

## References

1. Hazan J, Bester AC. CRISPR-Based Approaches for the High-Throughput Characterization of Long Non-Coding RNAs. Noncoding RNA [Internet]. 2021;7. Available from: http://www.ncbi.nlm.nih.gov/pubmed/34940760

2. Mattick JS, Amaral PP, Carninci P, Carpenter S, Chang HY, Chen LL, et al. Long non-coding RNAs: definitions, functions, challenges and recommendations. Nat Rev Mol Cell Biol [Internet]. 2023 [cited 2023 Apr 13];17:1–17. Available from: https://www.nature.com/articles/s41580-022-00566-8

3. Camilleri-Robles C, Amador R, Klein CC, Guigó R, Corominas M, Ruiz-Romero M. Genomic and functional conservation of lncRNAs: lessons from flies [Internet]. Mammalian Genome. Springer; 2022 [cited 2023 Apr 18]. p. 328–42. Available from: https://link.springer.com/article/10.1007/s00335-021-09939-4

4. Zhao Y, Li H, Fang S, Kang Y, Wu W, Hao Y, et al. NONCODE 2016: An informative and valuable data source of long non-coding RNAs. Nucleic Acids Res [Internet]. 2016 [cited 2020 Apr 6];44:D203–8. Available from: http://www.bioinfo.org/noncode/

5. Ulitsky I. Evolution to the rescue: Using comparative genomics to understand long non-coding RNAs. Nat Rev Genet. 2016;17:601–14.

6. Gao F, Cai Y, Kapranov P, Xu D. Reverse-genetics studies of lncRNAs-what we have learnt and paths forward [Internet]. Genome Biol. BioMed Central Ltd.; 2020 [cited 2021 Apr 1]. p. 1–23. Available from: 10.1186/s13059-020-01994-5

7. Ramilowski JA, Yip CW, Agrawal S, Chang JC, Ciani Y, Kulakovskiy I V., et al. Functional annotation of human long noncoding RNAs via molecular phenotyping. Genome Res [Internet]. 2020 [cited 2021 Apr 2];30:1060–72. Available from: http://www.genome.org/cgi/doi/10.http://creativecommons.org/licenses/by/4.0/www.genome.org

8. Nötzold L, Frank L, Gandhi M, Polycarpou-Schwarz M, Groß M, Gunkel M, et al. The long non-coding RNA LINC00152 is essential for cell cycle progression through mitosis in HeLa cells. Sci Rep. 2017;7:1–13.

9. Korkmaz G, Lopes R, Ugalde AP, Nevedomskaya E, Han R, Myacheva K, et al. Functional genetic screens for enhancer elements in the human genome using CRISPR-Cas9. Nat Biotechnol. 2016;34:192–8.

10. Liu Y, Cao Z, Wang Y, Guo Y, Xu P, Yuan P, et al. Genome-wide screening for functional long noncoding RNAs in human cells by Cas9 targeting of splice sites. Nat Biotechnol [Internet]. 2018 [cited 2021 Apr 1];36:1203–10. Available from: https://www.nature.com/articles/nbt.4283

11. Zhu S, Li W, Liu J, Chen CH, Liao Q, Xu P, et al. Genome-scale deletion screening of human long non-coding RNAs using a paired-guide RNA CRISPR-Cas9 library. Nat Biotechnol. 2016;34:1279–86.

12. Tao M, Mu Q, Zhang Y, Xie Z. Construction of a CRISPR-based paired-sgRNA library for chromosomal deletion of long non-coding RNAs. Quantitative Biology [Internet]. 2020 [cited 2021 Apr 1];8:31–42. Available from: 10.1007/s40484-020-0194-5

13. Fulco CP, Munschauer M, Anyoha R, Munson G, Grossman SR, Perez EM, et al. Systematic mapping of functional enhancer-promoter connections with CRISPR interference. Science (1979) [Internet]. 2016;354:769–73. Available from: http://www.sciencemag.org/lookup/doi/10.1126/science.aag2445

14. Joung J, Engreitz JM, Konermann S, Abudayyeh OO, Verdine VK, Aguet F, et al. Genome-scale activation screen identifies a lncRNA locus regulating a gene neighbourhood. Nature [Internet]. 2017;548:343–6. Available from: http://www.ncbi.nlm.nih.gov/pubmed/28792927

15. Bester AC, Lee JD, Chavez A, Lee YR, Nachmani D, Vora S, et al. An Integrated Genome-wide CRISPRa Approach to Functionalize lncRNAs in Drug Resistance. Cell [Internet]. 2018;173:649–664.e20. Available from: 10.1016/j.cell.2018.03.052

16. Xu D, Cai Y, Tang L, Han X, Gao F, Cao H, et al. A CRISPR/Cas13-based approach demonstrates biological relevance of vlinc class of long non-coding RNAs in anticancer drug response. Sci Rep [Internet]. 2020 [cited 2021 Apr 1];10:1–13. Available from: www.nature.com/scientificreports

17. Carlevaro-Fita J, Lanzós A, Feuerbach L, Hong C, Mas-Ponte D, Pedersen JS, et al. Cancer LncRNA Census reveals evidence for deep functional conservation of long noncoding RNAs in tumorigenesis. Commun Biol [Internet]. 2020;3:56. Available from: http://www.nature.com/articles/s42003-019-0741-7

18. Kirk JM, Kim SO, Inoue K, Smola MJ, Lee DM, Schertzer MD, et al. Functional classification of long non-coding RNAs by k-mer content. Nat Genet [Internet]. 2018;50:1474–82. Available from: http://www.ncbi.nlm.nih.gov/pubmed/30224646

19. Ehsani R, Drabløs F. Measures of co-expression for improved function prediction of long non-coding RNAs. BMC Bioinformatics. 2018;19:1–12.

20. Pyfrom SC, Luo H, Payton JE. PLAIDOH: A novel method for functional prediction of long non-coding RNAs identifies cancer-specific LncRNA activities. BMC Genomics. 2019;20:1–24.

21. Fernández M, Miranda-Saavedra D. Genome-wide enhancer prediction from epigenetic signatures using genetic algorithm-optimized support vector machines. Nucleic Acids Res [Internet]. 2012 [cited 2022 Apr 13];40:e77–e77. Available from: https://academic.oup.com/nar/article/40/10/e77/2411809

22. Wen J, Liu Y, Shi Y, Huang H, Deng B, Xiao X. A classification model for lncRNA and mRNA based on k-mers and a convolutional neural network. BMC Bioinformatics [Internet]. 2019;20:469. Available from: http://www.ncbi.nlm.nih.gov/pubmed/31519146

23. Zhang J, Zhang Z, Wang Z, Liu Y, Deng L. Ontological function annotation of long non-coding RNAs through hierarchical multi-label classification. Bioinformatics. 2018;34:1750–7.

24. Liu SJ, Horlbeck MA, Cho SW, Birk HS, Malatesta M, He D, et al. CRISPRi-based genome-scale identification of functional long noncoding RNA loci in human cells. Science [Internet]. 2017;355. Available from: http://www.ncbi.nlm.nih.gov/pubmed/27980086

25. Liu SJ, Malatesta M, Lien B V., Saha P, Thombare SS, Hong SJ, et al. CRISPRi-based radiation modifier screen identifies long non-coding RNA therapeutic targets in glioma. Genome Biol [Internet]. 2020 [cited 2021 Mar 18];21. Available from: 10.1186/s13059-020-01995-4

26. Haswell JR, Mattioli K, Gerhardinger C, Maass PG, Foster DJ, Fernandez PP, et al. Genome-Wide CRISPR Interference Screen Identifies Long Non-Coding RNA Loci Required for Differentiation and Pluripotency. SSRN Electronic Journal [Internet]. 2021 [cited 2021 Apr 1];2021.02.08.430256. Available from: 10.1101/2021.02.08.430256

27. Dao LTM, Galindo-Albarrán AO, Castro-Mondragon JA, Andrieu-Soler C, Medina-Rivera A, Souaid C, et al. Genome-wide characterization of mammalian promoters with distal enhancer functions. Nat Genet [Internet]. 2017 [cited 2023 Apr 19];49:1073–81. Available from: https://www.nature.com/articles/ng.3884

28. Chen T, Guestrin C. XGBoost: A scalable tree boosting system. Proceedings of the ACM SIGKDD International Conference on Knowledge Discovery and Data Mining [Internet]. New York, NY, USA: ACM; 2016 [cited 2023 Apr 19]. p. 785–94. Available from: 10.1145/2939672.2939785

29. Shapley LS. A value for n-person games. Contribution to the Theory of Games. 1953;

30. Gilbert LA, Larson MH, Morsut L, Liu Z, Brar GA, Torres SE, et al. CRISPR-mediated modular RNA-guided regulation of transcription in eukaryotes. Cell. 2013;154:442.

31. Horlbeck MA, Gilbert LA, Villalta JE, Adamson B, Pak RA, Chen Y, et al. Compact and highly active next-generation libraries for CRISPR-mediated gene repression and activation. Elife. 2016;5.

32. Chavez A, Scheiman J, Vora S, Pruitt BW, Tuttle M, P R Iyer E, et al. Highly efficient Cas9-mediated transcriptional programming. Nat Methods [Internet]. 2015 [cited 2021 Nov 26];12:326–8. Available from: https://www.nature.com/articles/nmeth.3312

33. Cao XM, Luo XG, Liang JH, Zhang C, Meng XP, Guo DW. Critical selection of internal control genes for quantitative real-time RT-PCR studies in lipopolysaccharide-stimulated human THP-1 and K562 cells. Biochem Biophys Res Commun. 2012;427:366–72.

34. Eekels JJM, Pasternak AO, Schut AM, Geerts D, Jeeninga RE, Berkhout B. A competitive cell growth assay for the detection of subtle effects of gene transduction on cell proliferation. Gene Ther [Internet]. 2012 [cited 2022 Apr 21];19:1058–64. Available from: https://www.nature.com/articles/gt2011191

35. Levin M, Stark M, Ofran Y, Assaraf YG. Deciphering molecular mechanisms underlying chemoresistance in relapsed AML patients: towards precision medicine overcoming drug resistance. Cancer Cell Int [Internet]. 2021;21:53. Available from: 10.1186/s12935-021-01746-w

36. Hashimshony T, Senderovich N, Avital G, Klochendler A, de Leeuw Y, Anavy L, et al. CEL-Seq2: Sensitive highly-multiplexed single-cell RNA-Seq. Genome Biol [Internet]. 2016 [cited 2023 Apr 10];17:1–7. Available from: https://genomebiology.biomedcentral.com/articles/10.1186/s13059-016-0938-8

37. Afgan E, Nekrutenko A, Grüning BA, Blankenberg D, Goecks J, Schatz MC, et al. The Galaxy platform for accessible, reproducible and collaborative biomedical analyses: 2022 update. Nucleic Acids Res [Internet]. 2022 [cited 2023 Apr 3];50:W345–51. Available from: https://academic.oup.com/nar/article/50/W1/W345/6572001

38. Blankenberg D, Gordon A, Von Kuster G, Coraor N, Taylor J, Nekrutenko A, et al. Manipulation of FASTQ data with galaxy. Bioinformatics [Internet]. 2010 [cited 2023 Apr 3];26:1783–5. Available from: https://academic.oup.com/bioinformatics/article/26/14/1783/178709

39. Kim D, Langmead B, Salzberg SL. HISAT: A fast spliced aligner with low memory requirements. Nat Methods [Internet]. 2015 [cited 2023 Apr 3];12:357–60. Available from: https://www.nature.com/articles/nmeth.3317

40. Liao Y, Smyth GK, Shi W. FeatureCounts: An efficient general purpose program for assigning sequence reads to genomic features. Bioinformatics [Internet]. 2014 [cited 2023 Apr 3];30:923–30. Available from: https://academic.oup.com/bioinformatics/article/30/7/923/232889

41. Tarazona S, García-Alcalde F, Dopazo J, Ferrer A, Conesa A. Differential expression in RNA-seq: A matter of depth. Genome Res [Internet]. 2011 [cited 2023 Apr 10];21:2213–23. Available from: https://genome.cshlp.org/content/21/12/2213.full

42. Tarazona S, Furió-Tarí P, Turrà D, Di Pietro A, Nueda MJ, Ferrer A, et al. Data quality aware analysis of differential expression in RNA-seq with NOISeq R/Bioc package. Nucleic Acids Res [Internet]. 2015 [cited 2023 Apr 10];43:e140–e140. Available from: https://academic.oup.com/nar/article/43/21/e140/2468096

43. Xie Z, Bailey A, Kuleshov M V., Clarke DJB, Evangelista JE, Jenkins SL, et al. Gene Set Knowledge Discovery with Enrichr. Curr Protoc [Internet]. 2021 [cited 2023 Apr 9];1:e90. Available from: https://onlinelibrary.wiley.com/doi/full/10.1002/cpz1.90

44. Chen EY, Tan CM, Kou Y, Duan Q, Wang Z, Meirelles G V., et al. Enrichr: Interactive and collaborative HTML5 gene list enrichment analysis tool. BMC Bioinformatics [Internet]. 2013 [cited 2023 Apr 9];14. Available from: https://pubmed.ncbi.nlm.nih.gov/23586463/

45. Kuleshov M V., Jones MR, Rouillard AD, Fernandez NF, Duan Q, Wang Z, et al. Enrichr: a comprehensive gene set enrichment analysis web server 2016 update. Nucleic Acids Res [Internet]. 2016 [cited 2023 Apr 9];44:W90–7. Available from: https://pubmed.ncbi.nlm.nih.gov/27141961/

46. Keenan AB, Torre D, Lachmann A, Leong AK, Wojciechowicz ML, Utti V, et al. ChEA3: transcription factor enrichment analysis by orthogonal omics integration. Nucleic Acids Res. 2019;47:W212–24.

47. Gerstein MB, Kundaje A, Hariharan M, Landt SG, Yan KK, Cheng C, et al. Architecture of the human regulatory network derived from ENCODE data. Nature [Internet]. 2012 [cited 2023 May 21];489:91–100. Available from: https://www.nature.com/articles/nature11245

48. Wang J, Zhuang J, Iyer S, Lin XY, Whitfield TW, Greven MC, et al. Sequence features and chromatin structure around the genomic regions bound by 119 human transcription factors. Genome Res [Internet]. 2012 [cited 2023 May 21];22:1798–812. Available from: https://genome.cshlp.org/content/22/9/1798.full

49. Wang J, Zhuang J, Iyer S, Lin XY, Greven MC, Kim BH, et al. Factorbook.org: A Wiki-based database for transcription factor-binding data generated by the ENCODE consortium. Nucleic Acids Res [Internet]. 2013 [cited 2023 May 21];41:D171–6. Available from: https://academic.oup.com/nar/article/41/D1/D171/1069417

50. Sloan CA, Chan ET, Davidson JM, Malladi VS, Strattan JS, Hitz BC, et al. ENCODE data at the ENCODE portal. Nucleic Acids Res [Internet]. 2016 [cited 2023 May 21];44:D726–32. Available from: https://academic.oup.com/nar/article/44/D1/D726/2502655

51. Luo Y, Hitz BC, Gabdank I, Hilton JA, Kagda MS, Lam B, et al. New developments on the Encyclopedia of DNA Elements (ENCODE) data portal. Nucleic Acids Res [Internet]. 2020 [cited 2023 May 21];48:D882–9. Available from: https://pubmed.ncbi.nlm.nih.gov/31713622/

52. Dunham I, Kundaje A, Aldred SF, Collins PJ, Davis CA, Doyle F, et al. An integrated encyclopedia of DNA elements in the human genome. Nature [Internet]. 2012 [cited 2023 May 21];489:57–74. Available from: https://pubmed.ncbi.nlm.nih.gov/22955616/

53. Huang R, Zhao L, Chen H, Yin RH, Li CY, Zhan YQ, et al. Megakaryocytic differentiation of K562 cells induced by PMA reduced the activity of respiratory chain complex IV. PLoS One [Internet]. 2014 [cited 2023 Feb 28];9:e96246. Available from: https://journals.plos.org/plosone/article?id=10.1371/journal.pone.0096246

54. Ren JG, Seth P, Everett P, Clish CB, Sukhatme VP. Induction of Erythroid Differentiation in Human Erythroleukemia Cells by Depletion of Malic Enzyme 2. PLoS One [Internet]. 2010 [cited 2023 May 17];5:e12520. Available from: https://journals.plos.org/plosone/article?id=10.1371/journal.pone.0012520

55. Fanucchi S, Mhlanga MM. Enhancer-Derived lncRNAs Regulate Genome Architecture: Fact or Fiction? Trends in Genetics [Internet]. 2017;33:375–7. Available from: https://linkinghub.elsevier.com/retrieve/pii/S0168952517300471

56. Libbrecht MW, Noble WS. Machine learning applications in genetics and genomics. Nat Rev Genet [Internet]. 2015;16:321–32. Available from: http://www.ncbi.nlm.nih.gov/pubmed/25948244

57. Pang LR, Huang MX, Li H, Chen G, Zhong GP, Yao B, et al. LINC00707 accelerates the proliferation, migration and invasion of clear cell renal cell carcinoma. Eur Rev Med Pharmacol Sci. 2020;24:6616–22.

58. Constanty F, Shkumatava A. lncRNAs in development and differentiation: From sequence motifs to functional characterization. Development [Internet]. 2021;148:dev182741. Available from: http://dev.biologists.org/content/148/1/dev182741.abstract

59. Dey BK, Mueller AC, Dutta A. Long non-coding RNAs as emerging regulators of differentiation, development, and disease. Transcription. 2014;5:e944014.

60. Liu Z, Zhang Y, Han X, Li C, Yang X, Gao J, et al. Identifying Cancer-Related lncRNAs Based on a Convolutional Neural Network. Front Cell Dev Biol. 2020;8:1–7.

61. Shalem O, Sanjana NE, Zhang F. High-throughput functional genomics using CRISPR-Cas9. Nat Rev Genet. Nature Publishing Group; 2015. p. 299–311.

62. Evers B, Jastrzebski K, Heijmans JPM, Grernrum W, Beijersbergen RL, Bernards R. CRISPR knockout screening outperforms shRNA and CRISPRi in identifying essential genes. Nat Biotechnol [Internet]. 2016 [cited 2021 May 3];34:631–3. Available from: http://www.nature.com/

63. Franklin J. The elements of statistical learning: data mining, inference and prediction. The Mathematical Intelligencer [Internet]. 2005 [cited 2023 Apr 13];27:83–5. Available from: https://link.springer.com/article/10.1007/BF02985802

64. Altman N, Krzywinski M. The curse(s) of dimensionality. Nat Methods. Nature Publishing Group; 2018. p. 399–400.

65. Nuñez JK, Chen J, Pommier GC, Cogan JZ, Replogle JM, Adriaens C, et al. Genome-wide programmable transcriptional memory by CRISPR-based epigenome editing. Cell [Internet]. 2021 [cited 2021 Apr 27];184:2503–2519.e17. Available from: 10.1016/j.cell.2021.03.025

66. Dempster JM, Boyle I, Vazquez F, Root DE, Boehm JS, Hahn WC, et al. Chronos: a cell population dynamics model of CRISPR experiments that improves inference of gene fitness effects. Genome Biol [Internet]. 2021 [cited 2023 Apr 9];22:1–23. Available from: https://genomebiology.biomedcentral.com/articles/10.1186/s13059-021-02540-7

67. Cao C, Zhang T, Zhang D, Xie L, Zou X, Lei L, et al. The long non-coding RNA, SNHG6–003, functions as a competing endogenous RNA to promote the progression of hepatocellular carcinoma. Oncogene [Internet]. 2017 [cited 2023 Feb 28];36:1112–22. Available from: https://www.nature.com/articles/onc2016278

68. Chen K, Wang X, Wei B, Sun R, Wu C, Yang H ji. LncRNA SNHG6 promotes glycolysis reprogramming in hepatocellular carcinoma by stabilizing the BOP1 protein. Anim Cells Syst (Seoul) [Internet]. 2022 [cited 2023 Feb 28];26:369–79. Available from: https://www.tandfonline.com/doi/abs/10.1080/19768354.2022.2134206

69. Lu W, Cao F, Feng L, Song G, Chang Y, Chu Y, et al. LncRNA Snhg6 regulates the differentiation of MDSCs by regulating the ubiquitination of EZH2 [Internet]. J Hematol Oncol. BioMed Central Ltd; 2021 [cited 2023 Feb 28]. p. 1–4. Available from: https://jhoonline.biomedcentral.com/articles/10.1186/s13045-021-01212-0

70. Wang HS, Zhang W, Zhu HL, Li QP, Miao L, Miao L. Long noncoding RNA SNHG6 mainly functions as a competing endogenous RNA in human tumors [Internet]. Cancer Cell Int. BioMed Central Ltd.; 2020 [cited 2023 Feb 28]. p. 1–10. Available from: https://cancerci.biomedcentral.com/articles/10.1186/s12935-020-01303-x

71. Liu F, Tian T, Zhang Z, Xie S, Yang J, Zhu L, et al. Long non-coding RNA SNHG6 couples cholesterol sensing with mTORC1 activation in hepatocellular carcinoma. Nat Metab [Internet]. 2022 [cited 2023 Feb 28];4:1022–40. Available from: https://www.nature.com/articles/s42255-022-00616-7

72. Xu M, Chen X, Lin K, Zeng K, Liu X, Xu X, et al. LncRNA SNHG6 regulates EZH2 expression by sponging miR-26a/b and miR-214 in colorectal cancer. J Hematol Oncol [Internet]. 2019 [cited 2023 Feb 28];12:1–17. Available from: https://link.springer.com/articles/10.1186/s13045-018-0690-5

73. Lan Z, Yao X, Sun K, Li A, Liu S, Wang X. The Interaction Between lncRNA SNHG6 and hnRNPA1 Contributes to the Growth of Colorectal Cancer by Enhancing Aerobic Glycolysis Through the Regulation of Alternative Splicing of PKM. Front Oncol. 2020;10:363.

74. Weng H, Huang H, Chen J. RNA N 6-Methyladenosine Modification in Normal and Malignant Hematopoiesis. Adv Exp Med Biol [Internet]. Springer New York LLC; 2019 [cited 2023 Apr 13]. p. 75–93. Available from: https://link.springer.com/chapter/10.1007/978-981-13-7342-8_4

75. Ferreira R, Ohneda K, Yamamoto M, Philipsen S. GATA1 Function, a Paradigm for Transcription Factors in Hematopoiesis. Mol Cell Biol [Internet]. 2005 [cited 2023 Apr 13];25:1215–27. Available from: https://journals.asm.org/doi/10.1128/MCB.25.4.1215-1227.2005

76. Zimta AA, Tomuleasa C, Sahnoune I, Calin GA, Berindan-Neagoe I. Long non-coding RNAs in myeloid malignancies. Front Oncol. 2019;9:1048.

77. Gil N, Ulitsky I. Regulation of gene expression by cis-acting long non-coding RNAs. Nat Rev Genet [Internet]. 2020;21:102–17. Available from: http://www.ncbi.nlm.nih.gov/pubmed/31729473

78. Zou Q, Du X, Zhou L, Yao D, Dong Y, Jin J. A short peptide encoded by long non-coding RNA small nucleolar RNA host gene 6 promotes cell migration and epithelial–mesenchymal transition by activating transforming growth factor-beta/SMAD signaling pathway in human endometrial cells. Journal of Obstetrics and Gynaecology Research [Internet]. 2023 [cited 2023 May 21];49:232–42. Available from: https://onlinelibrary.wiley.com/doi/full/10.1111/jog.15476

