## Supplementary Figures and Tables S1-S6 for "Integration of transcription regulation and functional genomic data reveals lncRNA SNHG6’s role in hematopoietic differentiation and leukemia"

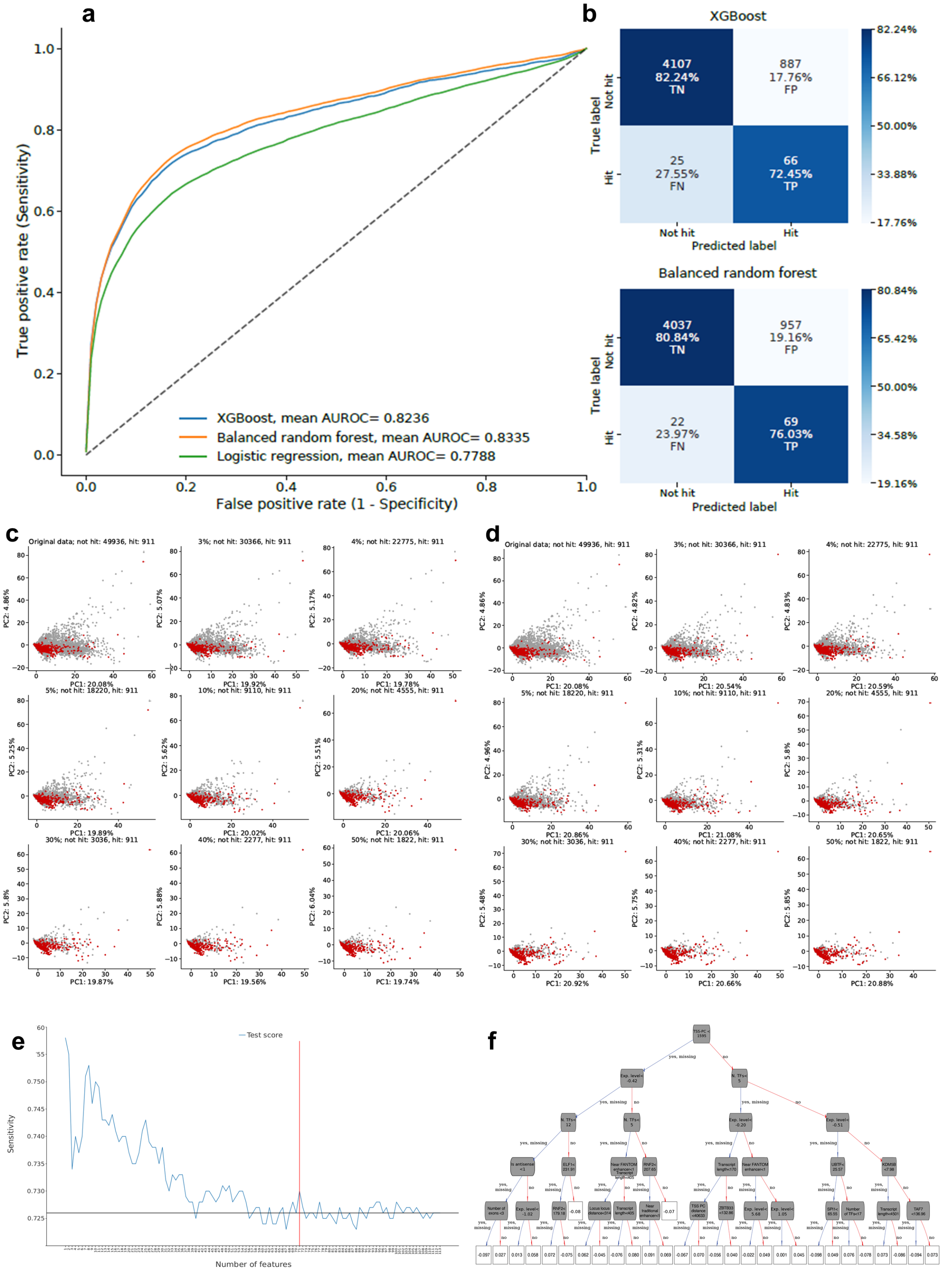
a
True positive rate (Sensitivity)
False positive rate (1 - Specificity)
XGBoost, mean AUROC= 0.8236
Balanced random forest, mean AUROC= 0.8335
Logistic regression, mean AUROC= 0.7788
b
XGBoost
4107
82.24%
TN
887
17.76%
FP
25
27.55%
FN
66
72.45%
TP
True label
Not hit
Hit
Predicted label
Balanced random forest
4037
80.84%
TN
957
19.16%
FP
22
23.97%
FN
69
76.03%
TP
c
Original data; not hit: 49936, hit: 911
3%; not hit: 30366, hit: 911
4%; not hit: 22775, hit: 911
5%; not hit: 18220, hit: 911
10%; not hit: 9110, hit: 911
20%; not hit: 4555, hit: 911
30%; not hit: 3036, hit: 911
40%; not hit: 2277, hit: 911
50%; not hit: 1822, hit: 911
d
e
Test score
Sensitivity
Number of features
f

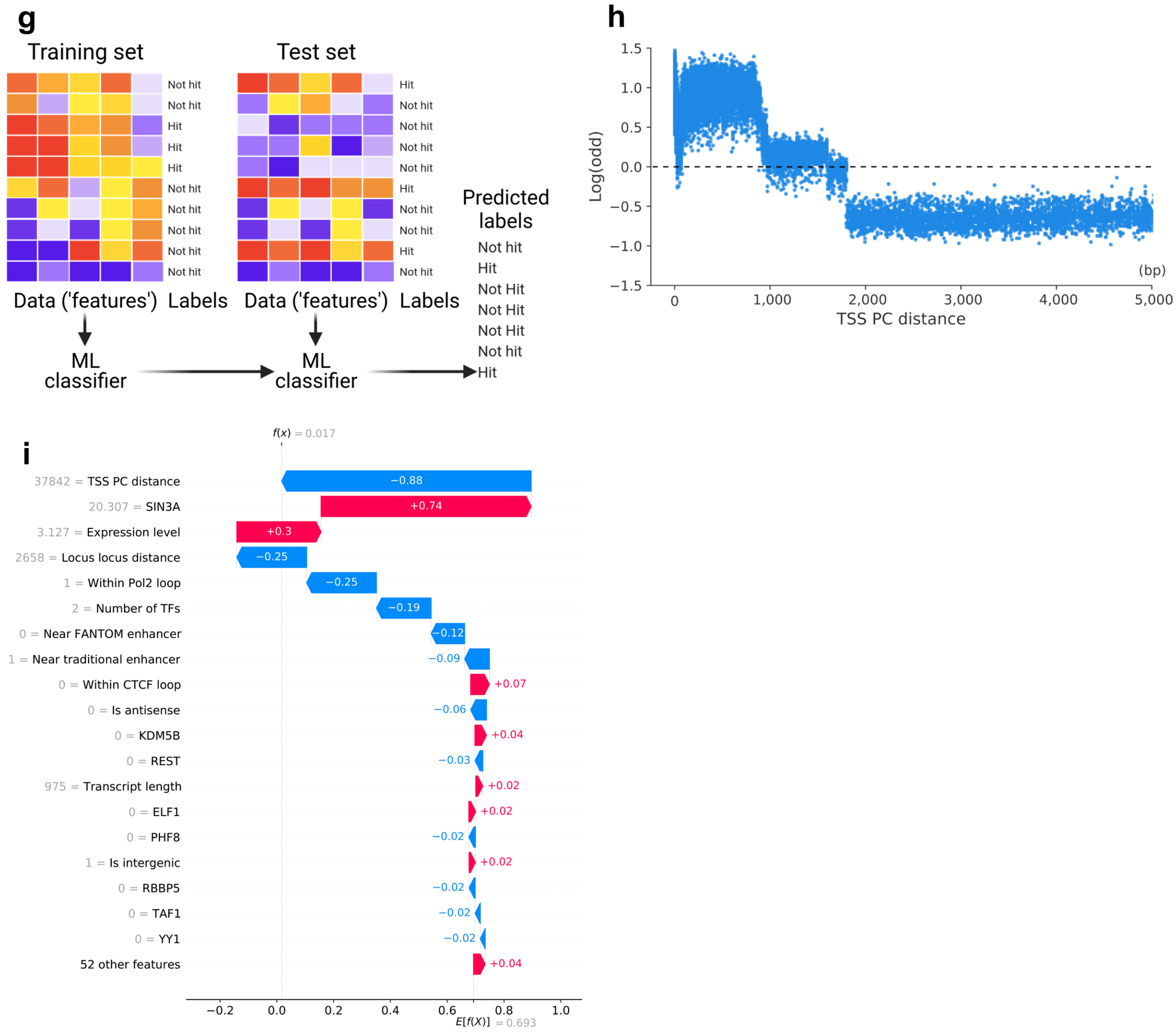


**Supplemental Fig. S1. Building the machine learning algorithm.** (a) ROC curves comparing XGBoost, balanced random forest, and logistic regression models. (b) XGBoost (upper) and balanced random forest (lower) confusion matrices. Models were trained on 90% of the data, and ROC curves and confusion matrices show predictive value on the remaining 10%. Percentages from confusion matrices are row-normalized. (c, d) Under-sampling PCA. PCA of random under-sampling of the majority class (i.e. not hit) without (c) and with (d) replacement, plotting the complete dataset (upper-left plot) plus 8 sampling strategies. PCA values based on 130 numeric features showing the removed not hit transcripts. Red dots: hit; grey dots: not hit. (e) Recursive feature elimination. Iteratively, one feature was removed to train a new model, removing the least important. Red and black lines denote the optimal number of features (n = 71) and sensitivity value using 143 features, respectively. (f) XGBoost first residual-tree. Tree nodes are represented as rounded grey boxes, and squared white boxes are the tree leafs. (g) Process followed for model training. The functional screening based on CRISPRi and the ENCODE Transcription Factor datasets were split into 90% for the training set (adopting a stratified cross-validation) and 10% for the testing set; along with binary labels indicating whether the lncRNA locus is either a hit or not hit. (h) SHAP dependence plot for the TSS PC distance feature. Each blue dot denotes a lncRNA. Positive odd values (above dashed line) contribute towards prediction of hits. (i) Detailed explanation for the INFLAMeR score of SNHG6. The final INFLAMeR score of SNHG6 was 0.504.

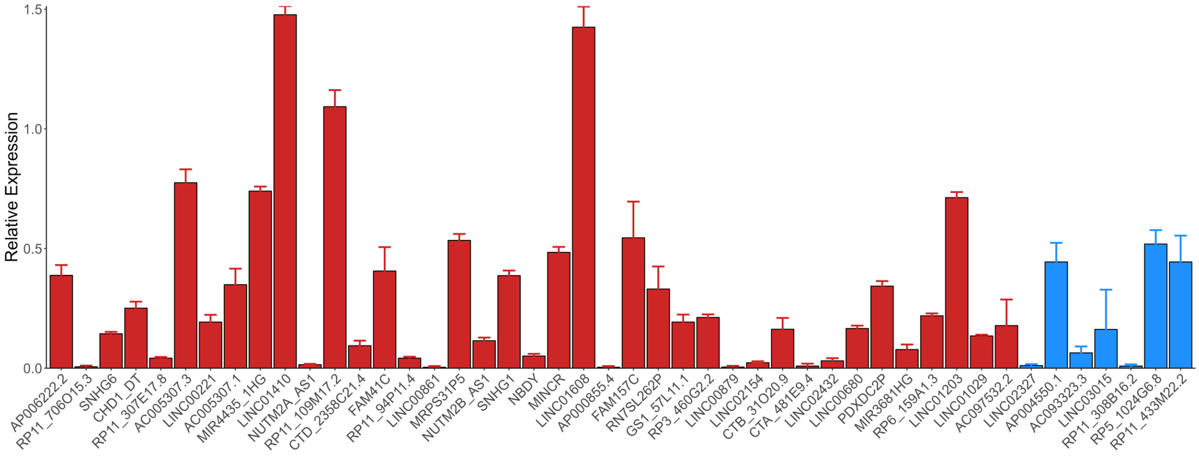


**Supplemental Fig. S2. Confirming sgRNA knockdown (KD) efficiency.** KD of thirty-nine lncRNAs predicted to be functional (red) and seven lncRNAs predicted to be non-functional (blue) was confirmed by qPCR. Expression is given relative to samples transduced with a non-targeting control sgRNA.

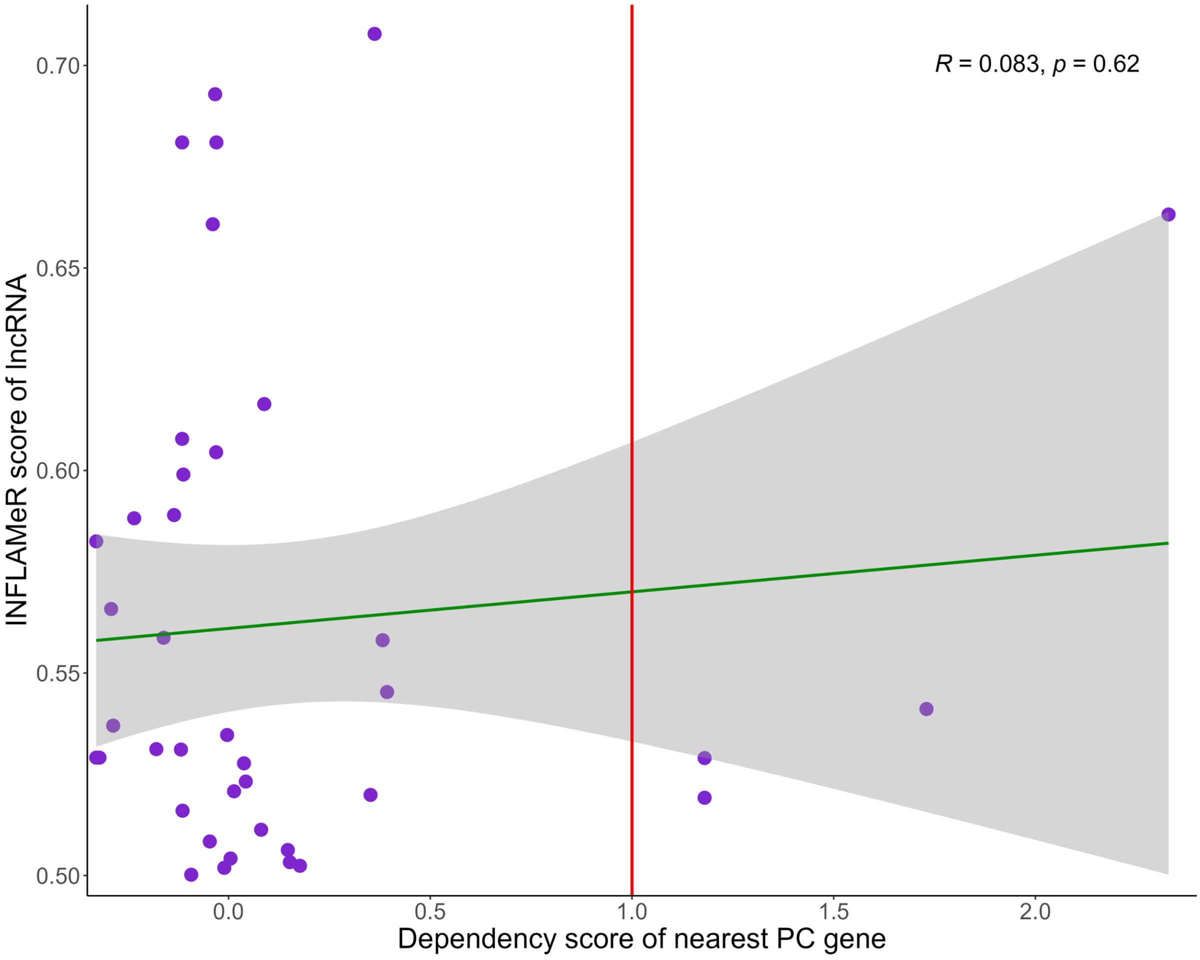


**Supplemental Fig. S3. There was no correlation between INFLAMeR score and the essentiality of neighboring protein-coding (PC) genes.** Genes with a dependency score > 1 (red line) are considered essential. Green line represents the linear regression with 95% confidence interval (grey).

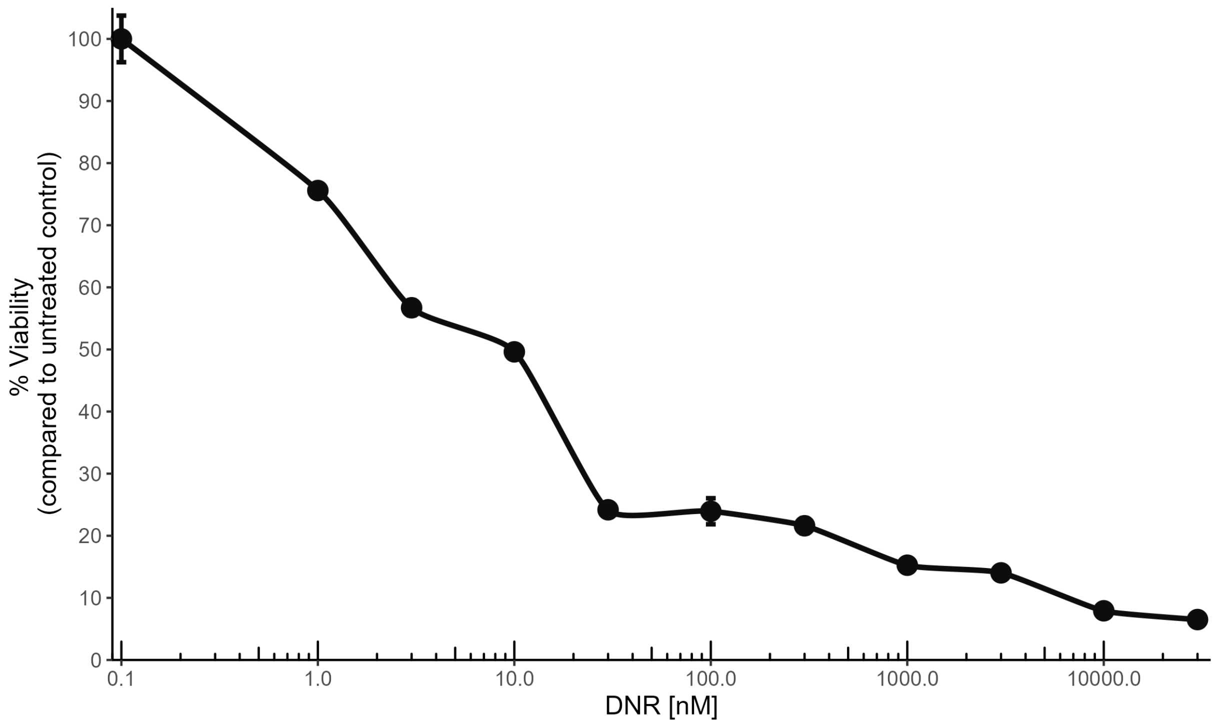


**Supplemental Fig. S4. Calibration of anticancer drug sensitivity assay.** K562 cells were incubated with the indicated concentrations of daunorubicin (DNR) for 72 h and their viability was determined by flow cytometry relative to that of untreated cells. Values are given as the mean ± SD for n = 3 biological replicates.

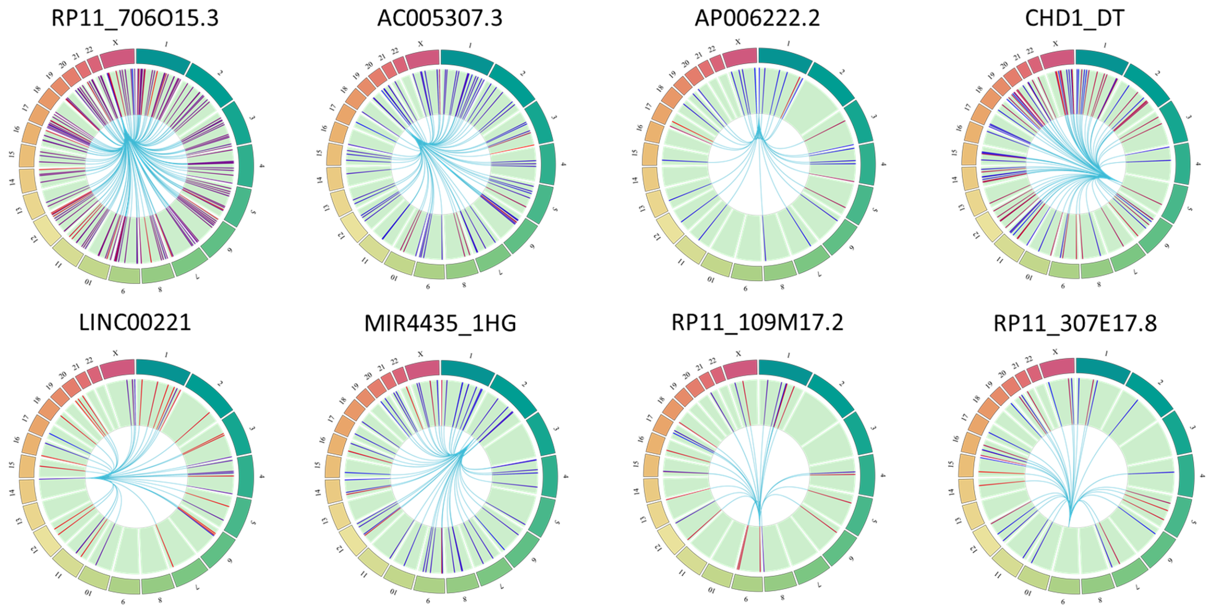


**Supplemental Fig. S5. KD of the indicated lncRNAs affected the expression of genes across the genome.** Red represents upregulation, blue represents downregulation.

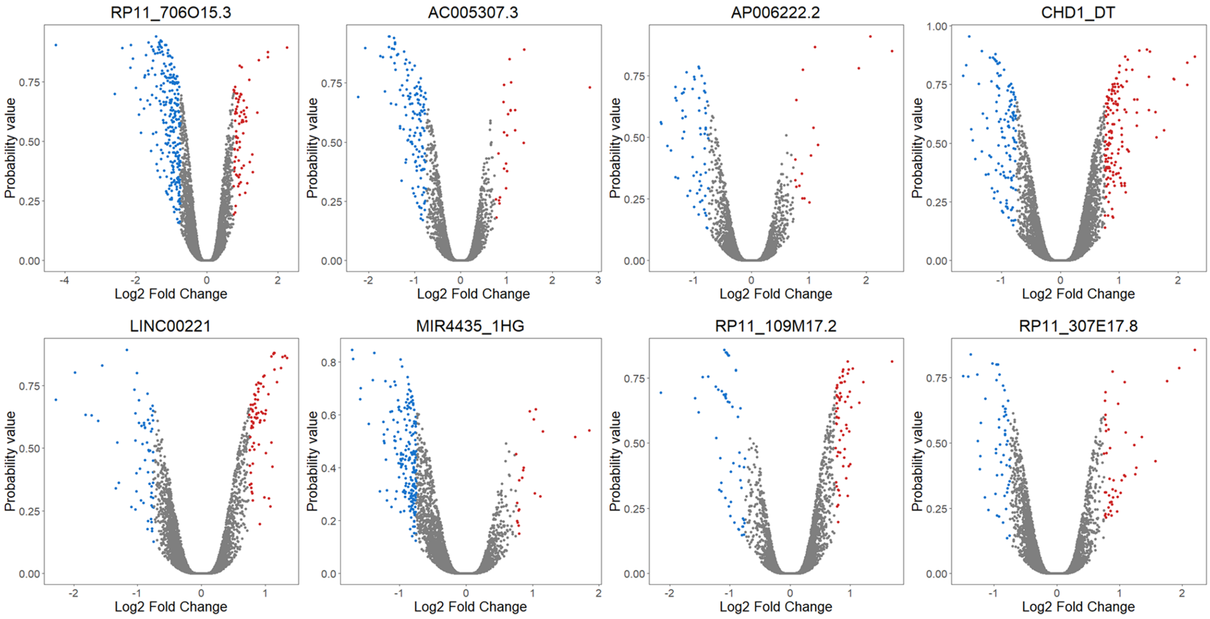


**Supplemental Fig. S6. Differentially expressed genes from each sample.** KD of the indicated lncRNAs generally led to a higher proportion of downregulated genes (blue) compared to upregulated genes (red).

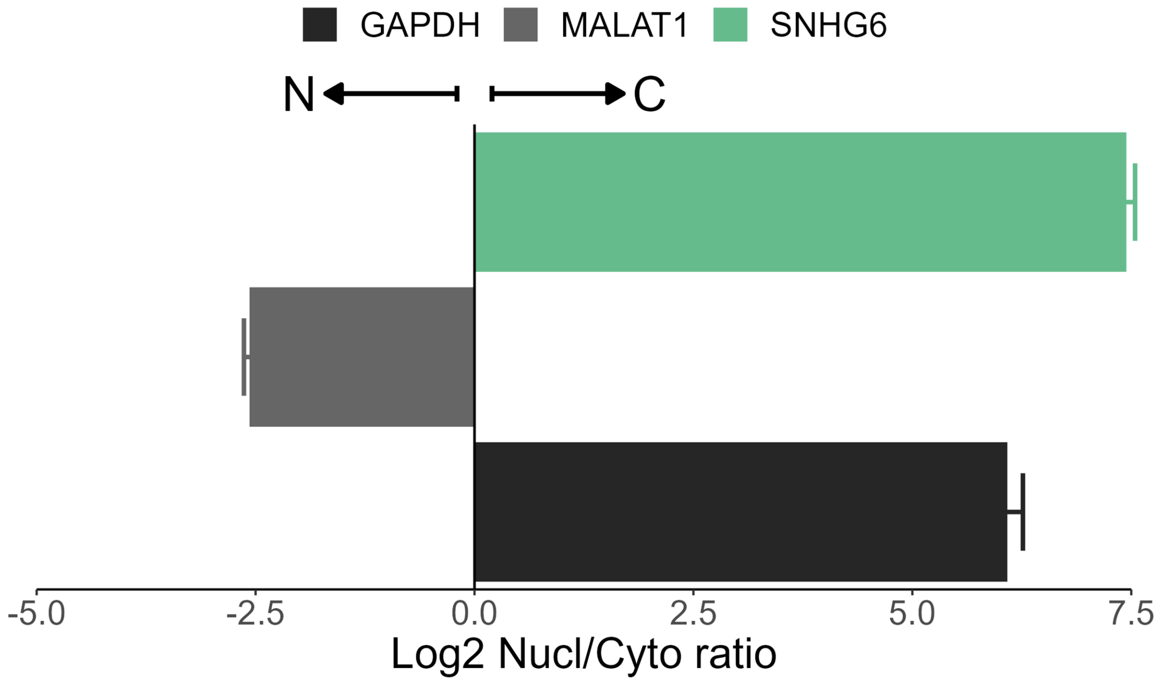


**Supplemental Fig. S7. SNHG6 subcellular localization.** The enrichment of SNHG6 was measured by qPCR in the nuclear/cytoplasmic fractions of K562 cells. MALAT1 and GAPDH were used as nuclear and cytoplasmic controls, respectively.

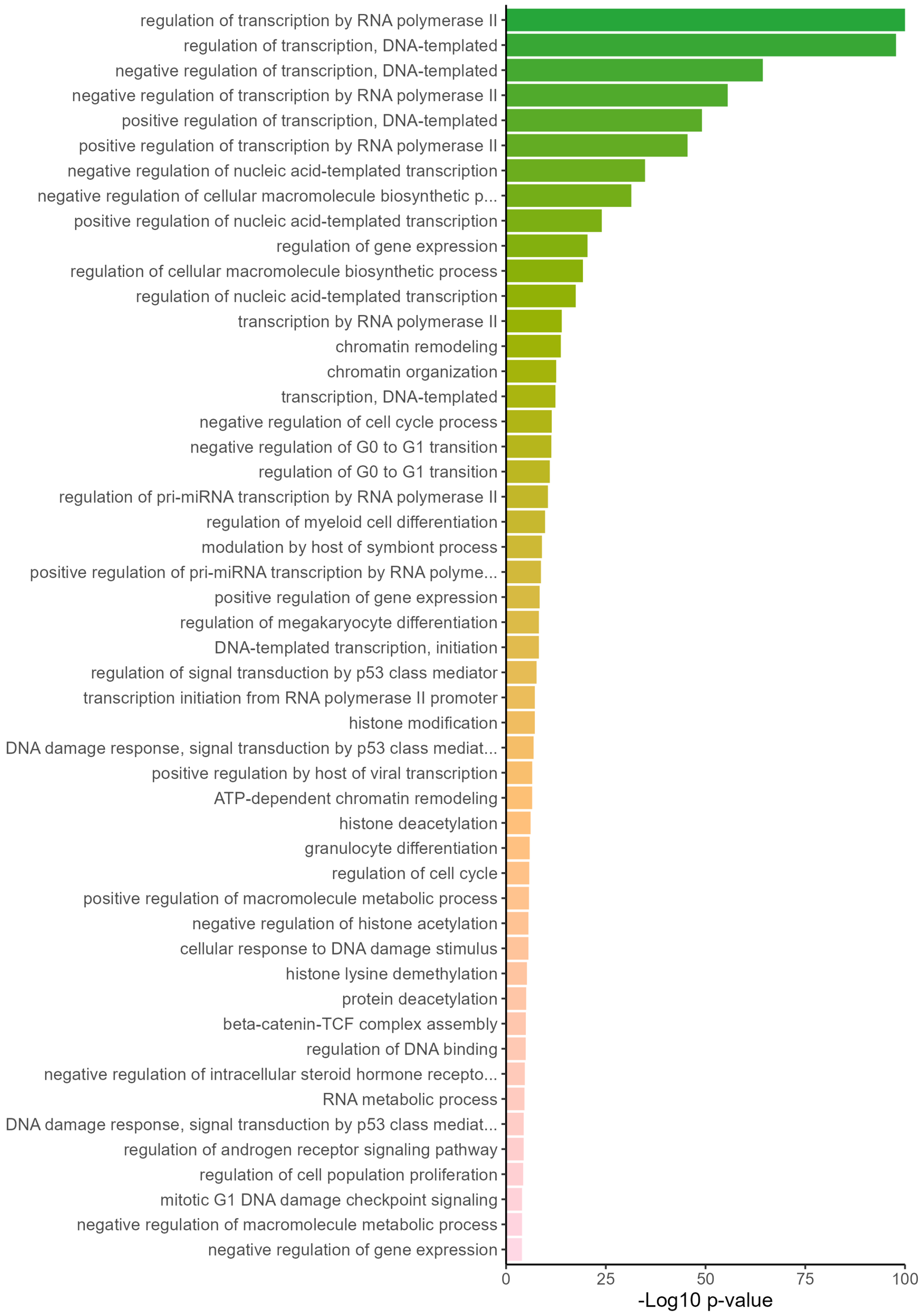


**Supplemental Fig. S8. Gene ontology analysis for the transcription factors that bind the promoter of SNHG6 in K562.** The top 50 pathways are shown.

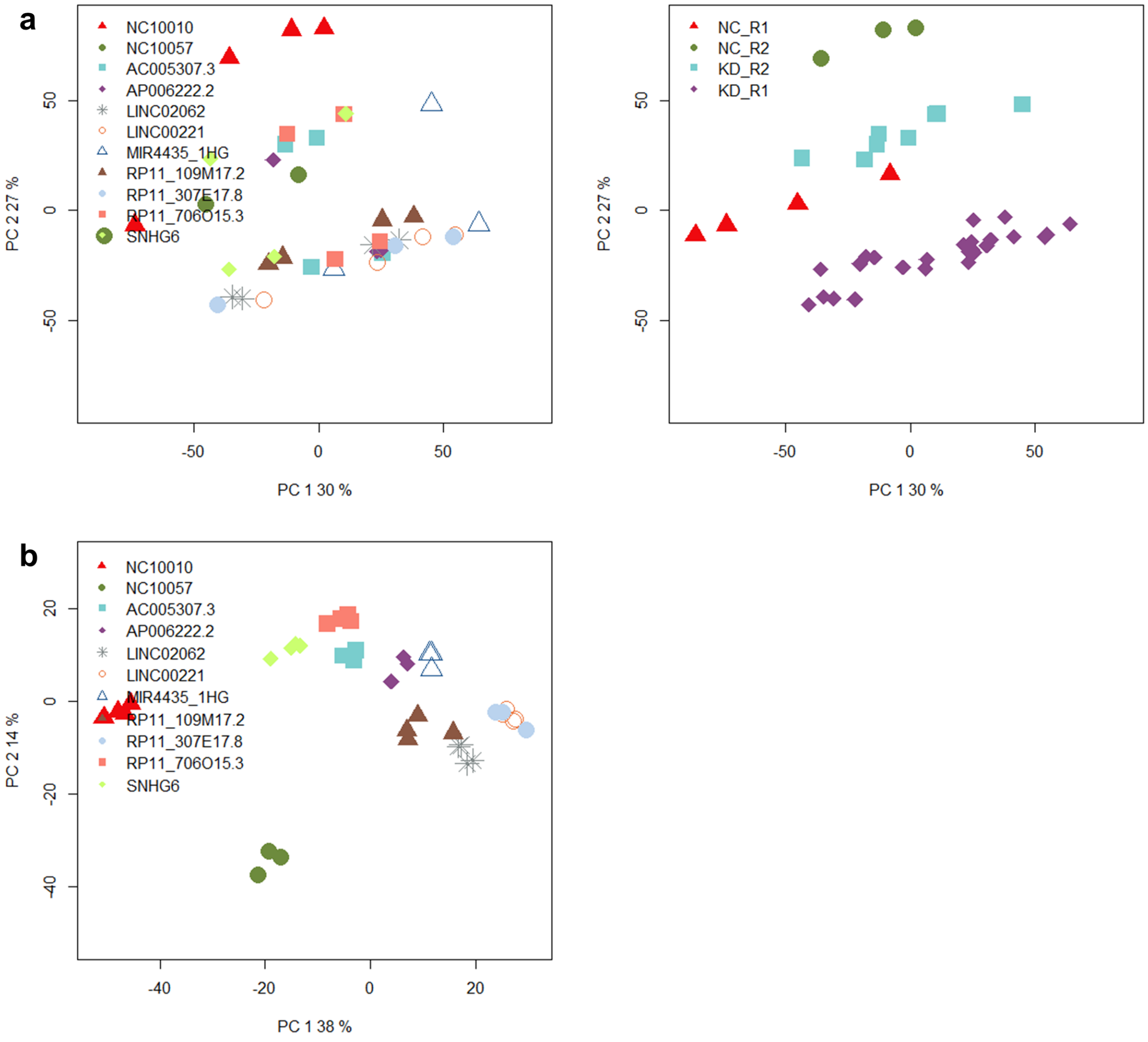


**Supplemental Fig. S9. Compensating for batch effect from RNA-Seq due to technical issues caused by multiple sequencing runs.** (a) Prior to batch effect correction, the samples were clustered not based on the biological samples (left), but rather based on the sequencing run (right). NC_: sgRNA negative control (NC) samples. KD_: samples with lncRNA knockdown (KD). R1 and R2: first and second sequencing run, respectively. (b) After batch effect correction, the samples were clustered according to the target KD, but the sgRNA control samples were distinctly clustered from all lncRNA KD samples.

**Supplemental Table S1. The 143 features included in the initial algorithm.**

| | | | | | | | | |
|---|---|---|---|---|---|---|---|---|
| Log2 FPKM | Is intergenic | Is antisense | TSS PC distance | Locus-locus distance | Locus is heterozygous deleted | Locus is amplified | Near cancer associated SNP | Near fantom enhancer |
| Near hnisz super enhancer | Near hnisz enhancer | Within CTCF loop | Within Pol2 loop | Transcript length | Number of exons | Has mouse ortholog | Number of TFs | ATF1 |
| ATF2 | CBX3 | CEBPB | CHD2 | CREB1 | CTCF | E2F1 | E2F6 | EGR1 |
| ELF1 | EP300 | ESRRA | FOS | FOSL1 | FOXM1 | GABPA | GTF2F1 | HCFC1 |
| HDAC1 | IKZF1 | IRF1 | JUND | KDM5B | MAFF | MAFK | MAZ | MTA3 |
| MYC | NCOR1 | NFIC | NR2C2 | PHF8 | POLR2A | RAD21 | RBBP5 | REST |
| RNF2 | SAP30 | SIN3A | SMARCA4 | SP1 | SPI1 | SREBF2 | TAF1 | TAF7 |
| TBP | TEAD4 | THAP1 | UBTF | YY1 | ZBTB33 | ZBTB7A | ZNF384 | |
| *Locus is homozygous deleted* | *Near vista enhancer* | *ARID3A* | *ATF3* | *BACH1* | *BCLAF1* | *BHLHE40* | *BRCA1* | *CBX8* |
| *CEBPZ* | *CHD1* | *CHD7* | *CTBP2* | *CTCFL* | *CUX1* | *E2F4* | *ELK1* | *ETS1* |
| *EZH2* | *FOSL2* | *FOXA1* | *GATA1* | *GATA2* | *GATA3* | *HDAC2* | *HDAC6* | *HSF1* |
| *JUN* | *KDM1A* | *KDM5A* | *MAX* | *MEF2A* | *MXI1* | *MYBL2* | *NANOG* | *NFE2* |
| *NFYA* | *NFYB* | *NR2F2* | *NR3C1* | *NRF1* | *PML* | *POU5F1* | *RCOR1* | *RELA* |
| *RFX5* | *RXRA* | *SETDB1* | *SIX5* | *SMARCB1* | *SMARCC2* | *SMC3* | *SREBF1* | *SRF* |
| *STAT5A* | *SUPT20H* | *SUZ12* | *TAL1* | *TBL1XR1* | *TCF12* | *TCF7L2* | *TRIM28* | *USF1* |
| *USF2* | *ZC3H11A* | *ZKSCAN1* | *ZMIZ1* | *ZNF143* | *ZNF217* | *ZNF263* | *ZNF274* | *ZZZ3* |

*Italics*: not included in the final algorithm.

**Supplemental Table S2. Cost-sensitive model metrics.**

| **Model** | **Sensitivity** | **Specificity** | **AUROC** | **F1** | **Precision** | **Brier score** |
|---|---|---|---|---|---|---|
| XGBoost | 0.7245 | 0.8224 | 0.8236 | 0.1264 | 0.0693 | 0.1638 |
| Balanced random forest | 0.7603 | 0.8084 | 0.8335 | 0.1240 | 0.0675 | 0.1460 |
| Logistic regression | 0.6165 | 0.8569 | 0.7788 | 0.1304 | 0.0629 | 0.1442 |

Values based on the mean of three randomization seeds of the test set.

**Supplemental Table S3. Under-sampling strategies without replacement.**

| **Sampling strategy** | **Sensitivity** | **Specificity** | **AUROC** | **F1** | **Precision** |
|---|---|---|---|---|---|
| 3% | 0.1627 | 0.9961 | 0.8250 | 0.2360 | 0.4363 |
| 4% | 0.2001 | 0.9942 | 0.8270 | 0.2621 | 0.3868 |
| 5% | 0.2341 | 0.9924 | 0.8281 | 0.2818 | 0.3583 |
| 10% | 0.3556 | 0.9826 | 0.8270 | 0.3073 | 0.2716 |
| 20% | 0.4946 | 0.9588 | 0.8292 | 0.2633 | 0.1797 |
| 30% | 0.5638 | 0.9342 | 0.8302 | 0.2182 | 0.1354 |
| 40% | 0.6114 | 0.9111 | 0.8289 | 0.1888 | 0.1117 |
| 50% | 0.6458 | 0.8894 | 0.8269 | 0.1679 | 0.0966 |

Preprocessing sampling strategies applied before XGBoost training.

**Supplemental Table S4. Under-sampling strategies with replacement.**

| Sampling strategy | Sensitivity | Specificity | AUROC | F1 | Precision |
|---|---|---|---|---|---|
| 3% | 0.1895 | 0.9945 | 0.8238 | 0.2531 | 0.3858 |
| 4% | 0.2301 | 0.9928 | 0.8257 | 0.2815 | 0.3665 |
| 5% | 0.2594 | 0.9906 | 0.8258 | 0.2918 | 0.3356 |
| 10% | 0.3604 | 0.9807 | 0.8307 | 0.2980 | 0.2546 |
| 20% | 0.4873 | 0.9589 | 0.8316 | 0.2609 | 0.1784 |
| 30% | 0.5675 | 0.9337 | 0.8301 | 0.2183 | 0.1353 |
| 40% | 0.6172 | 0.918 | 0.8306 | 0.1915 | 0.1134 |
| 50% | 0.6312 | 0.8897 | 0.8253 | 0.1646 | 0.0947 |

Preprocessing sampling strategies applied before XGBoost training.

**Supplemental Table S5. Model performance comparison.**

| Model | Sensitivity | Specificity | AUROC | F1 | Precision | Brier score |
|---|---|---|---|---|---|---|
| Cost-sensitive XGBoost 71 features | 0.7292 | 0.8227 | 0.8250 | 0.1275 | 0.0698 | 0.1634 |
| Cost-sensitive XGBoost 143 features | 0.7245 | 0.8224 | 0.8236 | 0.1264 | 0.0693 | 0.1638 |
| Balanced random forest 143 features | 0.7603 | 0.8084 | 0.8335 | 0.1240 | 0.0675 | 0.1460 |
| Cost-sensitive logistic regression 143 features | 0.6165 | 0.8569 | 0.7788 | 0.1304 | 0.0729 | 0.1442 |
| Under-sampling XGBoost without replacement 143 features | 0.6484 | 0.8894 | 0.8267 | 0.1679 | 0.0966 | 0.0907 |
| Under-sampling XGBoost with replacement 143 features | 0.6312 | 0.8897 | 0.8249 | 0.1646 | 0.0947 | 0.0915 |

Metrics based on the mean of 3 randomization seeds of the test set.

**Supplemental Table S6. Performance of 10-fold cross-validation (CV).**

| 10-fold CV | AUROC-1 | AUROC-2 | AUROC-3 |
|---|---|---|---|
| 1 | 0.78 | 0.80 | 0.80 |
| 2 | 0.82 | 0.85 | 0.84 |
| 3 | 0.83 | 0.82 | 0.82 |
| 4 | 0.83 | 0.80 | 0.82 |
| 5 | 0.79 | 0.85 | 0.82 |
| 6 | 0.83 | 0.82 | 0.83 |
| 7 | 0.82 | 0.85 | 0.85 |
| 8 | 0.83 | 0.82 | 0.82 |
| 9 | 0.84 | 0.85 | 0.84 |
| 10 | 0.87 | 0.84 | 0.81 |

Each column represents one randomization seed.
